# Piezo1 activation using Yoda1 inhibits macropinocytosis and proliferation of cancer cells

**DOI:** 10.1101/2021.05.14.444123

**Authors:** Masashi Kuriyama, Hisaaki Hirose, Toshihiro Masuda, Masachika Shudou, Jan Vincent V. Arafiles, Miki Imanishi, Masashi Maekawa, Yuji Hara, Shiroh Futaki

**Author notes:** School of Pharmaceutical Sciences, University of Shizuoka, Shizuoka 422-8526, Japan (Yuji Hara). These authors equally contributed to this work (Masashi Kuriyama, Hisaaki Hirose).

## Abstract

Macropinocytosis is a type of endocytosis accompanied by actin rearrangement-driven membrane deformation, such as lamellipodia formation and membrane ruffling, followed by macropinosome formation. A certain number of mammalian mechanosensors are sensitive to membrane deformation and tension. However, it remains unclear whether macropinocytosis is regulated by mechanosensors. Focusing on the mechanosensitive ion channel Piezo1, we found that Yoda1, a Piezo1 agonist, potently inhibits macropinocytosis induced by epidermal growth factor (EGF). Although studies with Piezo1 knockout cells suggest that Piezo1 itself is not physiologically indispensable for macropinocytosis regulation, Yoda1 inhibited ruffle formation depending on the extracellular Ca^2+^ influx through Piezo1 and on the activation of the calcium-activated potassium channel KCa3.1. This suggests that Ca^2+^ ions can regulate EGF-stimulated macropinocytosis. Moreover, Yoda1 impaired cancer cell proliferation, suggesting the impact of macropinocytosis inhibition. We propose the potential for cancer therapy by macropinocytosis inhibition through the regulation of a mechanosensitive channel activity.

## Introduction

Macropinocytosis is a large-scale endocytic pathway that accompanies membrane ruffling driven by actin rearrangement, followed by ruffle closure to form a vesicle called macropinosome (0.2 – 10 µm in diameter)^1–3^. Macropinosomes are significantly larger than vesicles produced by other endocytic pathways (∼ 100 nm in diameter), such as clathrin-mediated endocytosis, and macropinocytosis can non-selectively engulf a large volume of extracellular medium containing amino acids and proteins^3, 4^. Moreover, it has been shown that macropinocytosis plays a key role in promoting the survival of cancer cells through non-selective uptake of extracellular proteins and nutrients^5–7^, suggesting that the development of macropinocytosis inhibitors could be applicable for cancer therapy. From the viewpoint of the intracellular delivery research field, macropinocytosis has been applied and manipulated as an efficient internalization route for the uptake of macromolecules such as antibodies, proteins, and drugs^8, 9^. Therefore, understanding the molecular mechanisms of macropinocytosis and finding a new method to control this pathway should provide critical and useful insights not only for the development of cancer drugs, but also for drug delivery strategies.

The molecular mechanism of macropinocytosis has been studied in various types of cells, and there are two distinct forms of macropinocytosis: constitutive macropinocytosis and stimulated macropinocytosis^10^. Some specific cell types such as macrophages, dendritic cells, and Ras-transformed cancer cells constantly perform macropinocytosis without a stimulus. Recent studies have elucidated the molecular mechanisms underlying constitutive macropinocytosis. For instance, Canton *et al*. have reported that constitutive macropinocytosis in macrophages is dependent on calcium-sensing receptors (CaSR) and extracellular calcium ions (Ca^2+^)^11^. In contrast, macropinocytosis can be induced by various stimulants. For example, growth factors such as epidermal growth factor (EGF) and platelet-derived growth factor (PDGF), chemokines such as stromal cell-derived factor 1α (SDF1α, also known as CXCL12), and chemical compounds such as phorbol 12-myristate 13-acetate (PMA) can induce macropinocytosis in a wide range of cell types^12–15^. Notably, the molecular mechanisms by which EGF induces macropinocytosis have been extensively studied. EGF binds to the EGF receptor (EGFR) to activate small GTPases such as Rac1 and phosphatidylinositol 3-kinase (PI3K), leading to actin rearrangement and sequential conversion of phosphoinositides (*i.e.*, phosphorylation and dephosphorylation), followed by dynamic membrane ruffling, ruffle closure, and formation of macropinosomes^16^.

In addition to further progress in understanding growth factor-stimulated macropinocytosis^15^, a recent study reported that a decrease in plasma membrane tension could induce macropinocytosis^17^, implying that alteration in membrane tension may regulate macropinocytosis. Membrane tension has been suggested to significantly impact on various cellular functions^18^. A previous study showed that membrane tension regulates cell spreading and lamellipodial extension^19^. Eukaryotic cells have various mechanosensors that sense and respond to membrane tension^20, 21^. Mechanosensors respond to mechanical stimuli, such as alterations in membrane tension, to trigger signaling cascades that result in physiological responses^22^. To the best of our knowledge, there are no reports on the relationship between mechanosensors and macropinocytosis regulation.

Among mechanosensors, mechanosensitive ion channels that couple mechanical force with ion flux have been extensively studied to elucidate their physiological roles and gating mechanisms^21^. In particular, much attention has been paid to the recently identified Piezo1, a mammalian mechanosensitive ion channel^23^. Piezo1 is a non-selective cation channel for Na^+^, K^+^, and Ca^2+^, with a higher preference for Ca^2+^ over Na^+^ and K^+^ ions^23^. Piezo1 has been reported to play key roles in a plethora of physiological phenomena such as flow sensing in endothelial cells, bone formation, myotube formation, T cell activation and regulation of red blood cell volume^24–28^. Recent studies have suggested that Piezo1 is involved in the regulation of actin polymerization^29, 30^. In addition, Piezo1 is activated by mechanical stimuli as well as a small chemical compound called Yoda1, which was discovered by a high-throughput screen to activate Piezo1^31^. Yoda1 is a known Piezo1-specific agonist and is generally used to activate Piezo1^32^.

Since macropinocytosis is accompanied by membrane deformation and actin rearrangements, macropinocytosis may affect membrane tension and the activities of mechanosensitive channels or *vice versa*. The above studies and insights concerning Piezo1 motivated us to focus on the role of Piezo1 in EGF-stimulated macropinocytosis. In addition, it remains unclear whether Ca^2+^ ions affect the regulation of EGF-stimulated macropinocytosis. Therefore, we investigated the effect of Piezo1 activation using Yoda1 and extracellular Ca^2+^ influx on macropinocytosis. Here we show that Yoda1 potently inhibits EGF-stimulated macropinocytosis and that Yoda1 impairs cancer cell proliferation. This work suggests that Yoda1 enhances Ca^2+^ influx followed by aberrant activation of the calcium-activated potassium channel KCa3.1 and inhibition of Rac1 activation. We propose that Ca^2+^ is a of key factor in the regulation of EGF-stimulated macropinocytosis. Importantly, our study suggests the potential for cancer therapy by inhibiting macropinocytosis by regulating a mechanosensitive channel.

## Results

### Piezo1 activation using Yoda1 inhibits EGF-stimulated macropinocytosis in A431 cells

We used the human epidermoid carcinoma cell line A431, which expresses high levels of the EGFR and is a representative cell line for research on EGF-stimulated macropinocytosis. A previous study has reported that EGF-stimulated macropinocytosis in A431 cells can be evaluated by the amount of cellular uptake of tetramethylrhodamine (TMR)-conjugated dextran 70 kDa (TMR-dex70), a macropinosome marker, using flow cytometry and confocal microscopy analysis^33^.

To assess whether Piezo1 activation is involved in the regulation of EGF-stimulated macropinocytosis in A431 cells, we used Yoda1. A431 cells were incubated with TMR-dex70 in the presence or absence of EGF and Yoda1, and cellular uptake of TMR-dex70 was quantitatively evaluated by flow cytometry and observed by confocal microscopy. EGF treatment enhanced the cellular uptake of TMR-dex70, whereas Yoda1 significantly inhibited the EGF-induced uptake of TMR-dex70 in a concentration-dependent manner (**Figure 1A, B**), indicating that Yoda1 (1.5 µM) almost completely inhibited the cellular uptake of TMR-dex70 induced by EGF. On the other hand, Yoda1 did not substantively inhibit (∼10% inhibition at most) the clathrin-dependent cellular uptake of Alexa Fluor 568-labeled Transferrin (Tfn) into A431 cells (**Figure 1—figure supplement 1**). These results suggest that Piezo1 activation using Yoda1 selectively inhibits EGF-stimulated macropinocytosis.

**Figure 1.**
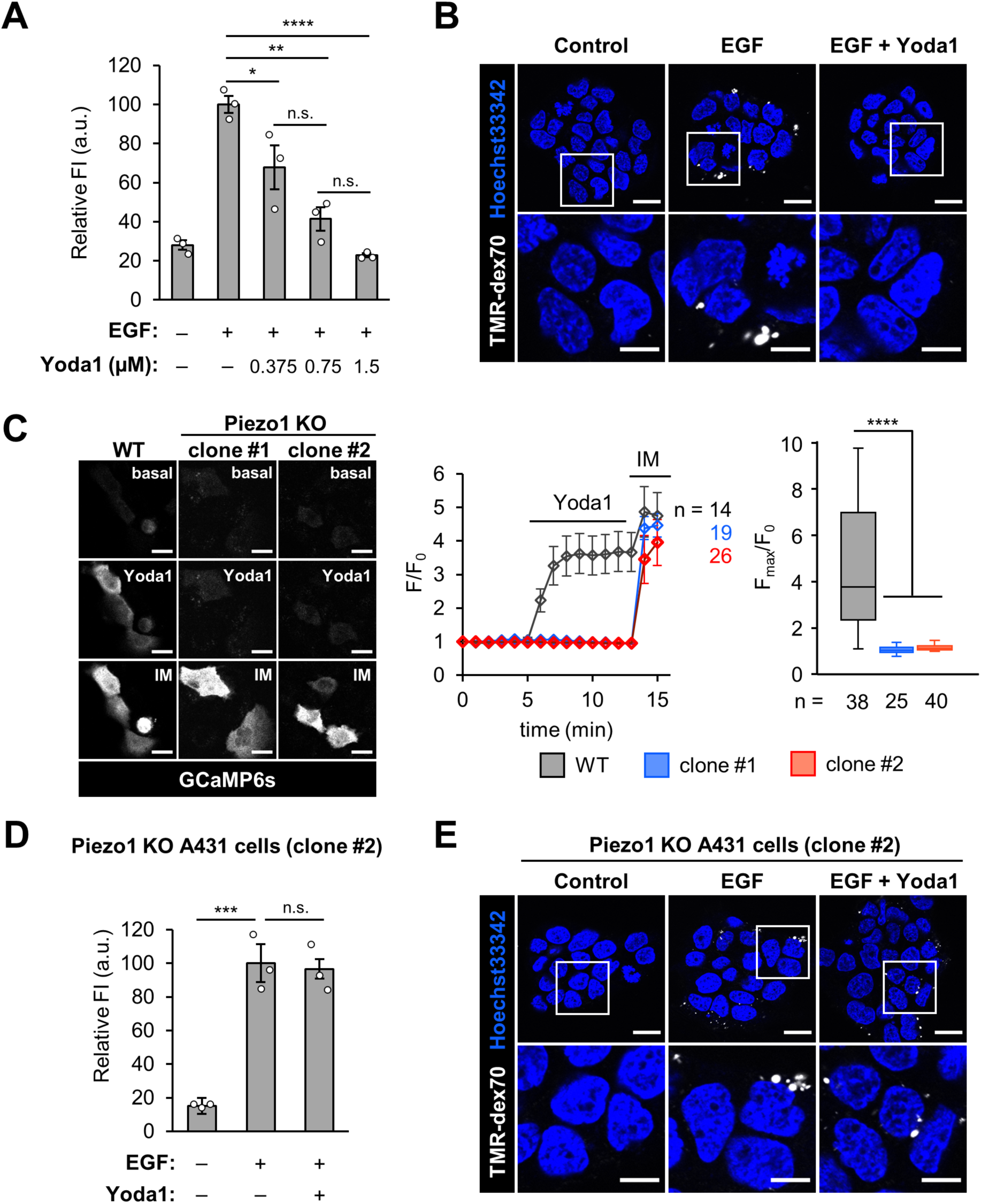
Piezo1 activation using Yoda1 inhibits macropinocytosis induced by EGF in A431 cells. (**A**) Flow cytometry analysis of TMR-dex70 uptake into A431 cells with or without EGF (20 nM) in the absence or presence of Yoda1 at the indicated concentration for 10 min. (**B**) Confocal microscopy observation of TMR-dex70 uptake into the cells stimulated with EGF (20 nM) in the absence or presence of Yoda1 (1.5 µM). The bottom images show enlarged views of the areas outlined by the white squares in the top images. (**C**) GCaMP6s fluorescence intensities in WT, Piezo1 KO clone#1 and clone #2 cells were recorded every 1 min. Yoda1 (1.5 µM) and ionomycin (5 µM) were added at 5 and 13 min after start of time-lapse imaging, respectively. Left: Representative images of GCaMP6s-expressing cells before and after addition of Yoda1 and ionomycin. Middle: Representative time-course of relative fluorescence intensity of GCaMP6s. Right: Quantification of maximum Yoda1-induced GCaMP6s intensity increase. Box and whiskers graph: line, median; box, upper and lower quartiles; whiskers, maxima and minima. (**D**) Flow cytometry analysis of TMR-dex70 uptake into Piezo1 KO A431 cells (clone #2) stimulated with EGF (20 nM) in the absence or presence of Yoda1 (1.5 µM) for 10 min. (**E**) Confocal microscopy observation of TMR-dex70 uptake into Piezo1 KO A431 cells treated as (**D**). The bottom images show enlarged views of the areas outlined by the white squares in the top images. Data represent the mean ± SEM (n = 3 independent biological replicates in (**A**) and (**D**); n = 14, 19, 26 cells for WT, clone #1 and #2, respectively) in (**C** middle)). Data represent in box plot (from left to right, n = 38, 25, 40 cells pooled from two independent experiments) in (**C** right). *, p < 0.05; **, p < 0.01; ***, p < 0.001; ****, p < 0.0001; n.s., not significant (one-way ANOVA followed by Tukey-Kramer’s post hoc test (**A**, **D**) or one-way ANOVA followed by Dunnett’s post hoc test (**C**)). Scale bars, (**B**, **C**, **E** top) 20 µm; (**E** bottom)10 µm. **Figure supplement 1.** Tfn uptake in the absence or presence of Yoda1. **Figure supplement 2.** Supporting data for characterization of Piezo1 knockout A431 cells. **Figure supplement 3.** Piezo1 gene expression examined by qPCR.

To determine whether the inhibitory effect of Yoda1 on macropinocytosis is Piezo1-dependent, we established Piezo1 knockout (KO) A431 cells, using the CRISPR/Cas9 system^34^. We obtained two clones (clones #1 and #2) in which Yoda1-induced Ca^2+^ influx, which can be detected by fluorescence of GCaMP6s, a genetically encoded Ca^2+^ indicator^35^, was completely abolished (**Figure 1C**). We also confirmed that ionomycin, a Ca^2+^ ionophore, induced Ca^2+^ influx in both wild-type and Piezo1 KO cells, indicating that Yoda1-induced Ca^2+^ is dependent on Piezo1 (**Figure 1C**). The DNA sequences around the CRISPR/Cas9-targeting region in these two clones were further confirmed by genome DNA sequencing, and all the detected alleles in clones #1 and #2 had CRISPR/Cas9-mediated gene mutations were as follows: clone #1 had 10, 16, and 33 bp-deletion mutations, and clone #2 had 9 and 10-bp deletion mutations (**Figure 1—figure supplement 2A**). Since there were alleles without frameshift mutations in both clones (*i.e.*, 33 and 9-bp deletions, resulting in 11 and 3-amino acid deletions of Piezo1 protein, referred to as Δ946-956 and Δ944-946, respectively), we investigated whether these two mutants lost the function of Piezo1. Piezo1 is a cation channel that allows Ca^2+^ influx^23^. Therefore, we monitored the fluorescence intensities of the Ca^2+^ indicator Fluo-8 in the absence or presence of Yoda1 in HEK293T cells transiently expressing full-length Piezo1, Δ946-956, or Δ944-946 (**Figure 1—figure supplement 2B**). The mutant Piezo1(Δ946-956) derived from the clone #1 allele slightly induced Ca^2+^ influx in the presence of Yoda1, whereas the other mutant Piezo1(Δ944-946) derived from the clone #2 allele did not. Therefore, we used clone #2 as Piezo1 KO A431 cells for the subsequent experiments. Piezo1 gene expression in A431 wild-type (WT) and Piezo1 KO cells was further confirmed by real-time quantitative PCR (qPCR) (**Figure 1—figure supplement 3**). The results indicated that the mRNA level of the Piezo1 coding region was decreased by over 90 % in the Piezo1 KO cell line, suggesting that there is minimal expression of Piezo1(Δ944-946) in the Piezo1 KO cells. Using the Piezo1 KO A431 cells, we conducted a dextran uptake assay. Flow cytometry analysis and confocal microscopy observation revealed that EGF induced macropinocytosis in Piezo1 KO A431 cells in both the absence and presence of Yoda1 (**Figure 1D, E**). These results show that Piezo1 itself is not physiologically required for EGF-stimulated macropinocytosis because macropinocytosis can be induced in Piezo1 KO A431 cells; however, the Piezo1 agonist Yoda1 inhibits macropinocytosis in a Piezo1-dependent manner.

**Figure 2.**
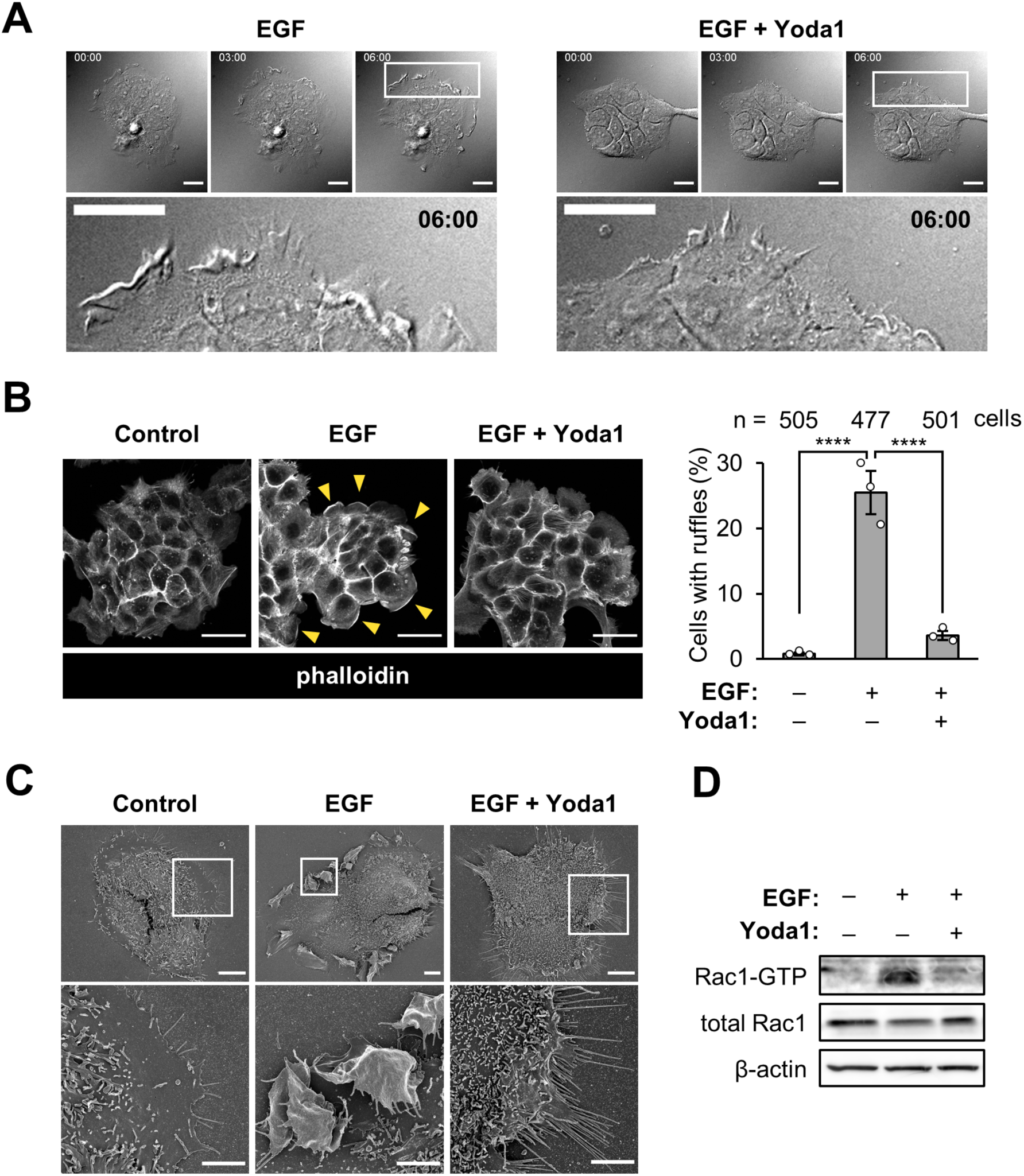
Piezo1 activation inhibits peripheral ruffle formation. (**A**) Live cell imaging of EGF- induced membrane ruffles in A431 cells. The cells were stimulated with EGF (20 nM) in the absence or presence of Yoda1 (1.5 µM). Live cell imaging was started immediately after adding EGF and Yoda1. Differential interference contrast (DIC) images at the indicated time points (0, 3 and 6 min) are shown. The bottom images show enlarged views of the areas outlined by the white squares in the images at 6 min. (**B**) F-actin staining with phalloidin. A431 cells were stimulated with EGF (20 nM) in the absence or presence of Yoda1 (1.5 µM) for 5 min, fixed, and stained with phalloidin-rhodamine to detect F-actin. Left: Representative images are shown. Yellow arrowheads indicate peripheral membrane ruffling area. Right: Quantification of cells with peripheral ruffles. Data represent the mean ± SEM (number of total counted cells pooled from three independent experiments: from left to right, 505, 477, and 501). (**C**) Scanning electron microscopy images of A431 cells stimulated with EGF (20 nM) in the absence or presence of Yoda1 (1.5 µM) for 5 min. The bottom images show enlarged views of the areas outlined by the white squares in the top images. (**D**) Rac1 activation pull-down assay. A431 cells treated with EGF (20 nM) and Yoda1 (1.5 μM) for 3 min were lysed and active Rac1 was precipitated using PAK-PBD beads. Active and total Rac1 protein were analyzed by performing SDS-PAGE followed by western blot. The representative images are shown. The experiments were performed twice with similar results. ****, P < 0.0001 (one-way ANOVA followed by Tukey-Kramer’s post hoc test (**B**)). Scale bars, (**A**) 20 µm; (**B**) 50 µm; (**C** top) 10 µm; (**C** bottom) 5 µm. **Figure supplement 1.** Yoda1 does not change intracellular pH. **Figure supplement 2.** Yoda1 does not inhibit phosphorylation of EGFR and Vav2. **Figure supplement 3.** Yoda1 does not affect cholesterol localization.

**Figure 3.**
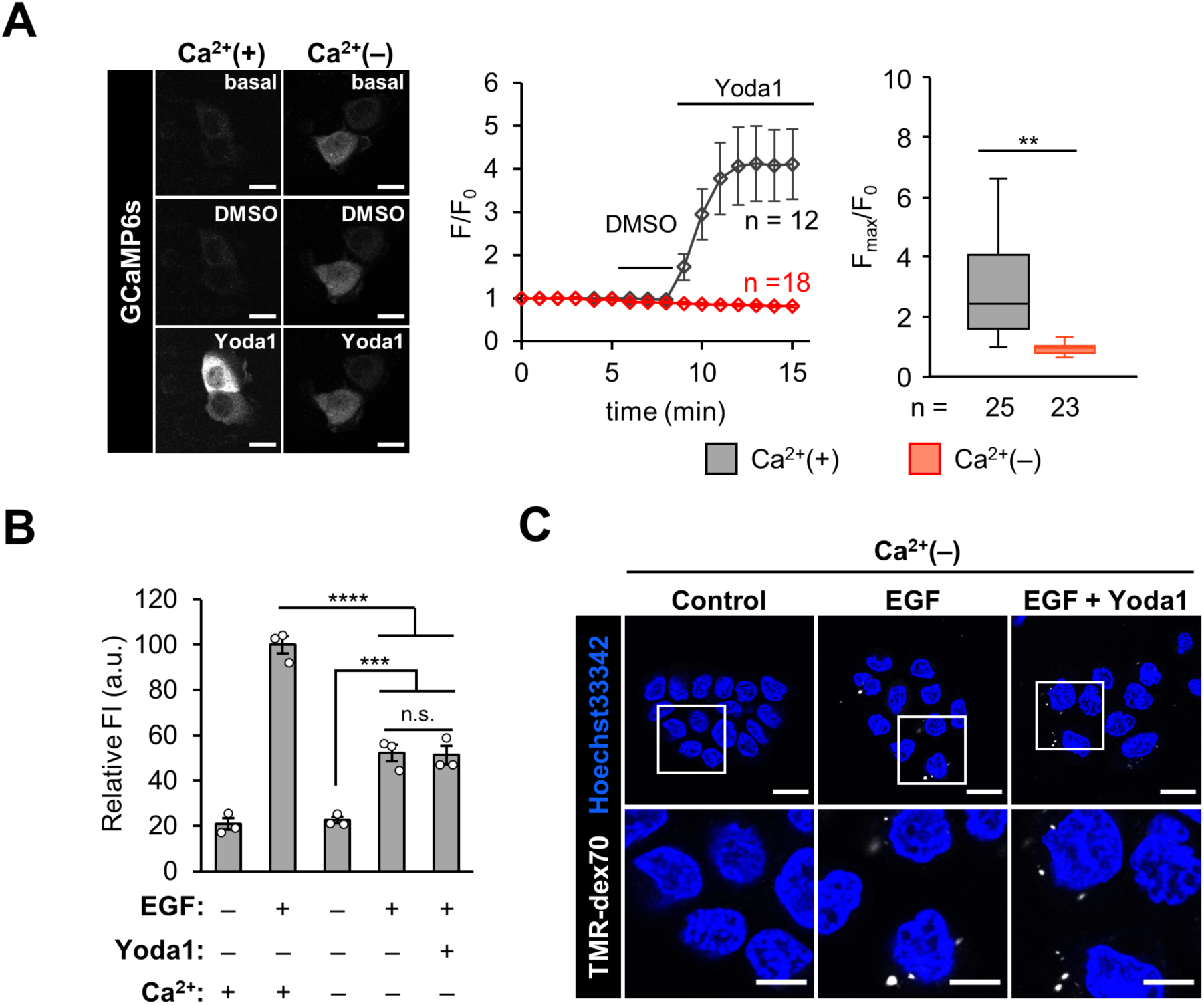
Macropinocytosis inhibition by Yoda1 requires extracellular Ca^2+^ influx. (**A**) A431 cells were transfected with GCaMP6s and then GCaMP6s fluorescence intensity was recorded every 1 min. DMSO and Yoda1 (1.5 µM) were added at 5 and 8 min after start of time-lapse imaging, respectively. Left: Representative images of the GCaMP6s-expressing cells treated DMSO and Yoda1 in the absence or presence of Ca^2+^ in culture media. Middle: Representative time-course of relative fluorescence intensity of GCaMP6s. Data represent the mean ± SEM (n = 12 and 18 cells for Ca^2+^(+) and Ca^2+^(–), respectively). Right: Quantification of maximum Yoda1-induced GCaMP6s intensity increase. Data represent in box plot (n = 25 and 23 cells, for Ca^2+^(+) and Ca^2+^(–), respectively, pooled from two independent experiments). Box and whiskers graph: line, median; box, upper and lower quartiles; whiskers, maxima and minima. (**B**) Flow cytometry analysis of TMR-dex70 uptake in Ca^2+^-free condition. The cells were stimulated with EGF (20 nM) for the uptake of TMR-dex70 in the absence or presence of Yoda1 (1.5 µM) and Ca²⁺ for 10 min. Data represent the mean ± SEM (n = 3 independent biological replicates). (**C**) Confocal microscopy observation of TMR-dex70 uptake in Ca^2+^-free condition. The cells were treated as (**B**). The bottom images show enlarged views of the areas outlined by the white squares in the top images. **, p < 0.01; ***, p < 0.001; ****, p < 0.0001; n.s., not significant (Student’s t-test (**A**) or one-way ANOVA followed by Tukey-Kramer’s post hoc test (**B**)). Scale bars, (**A**, **C** top) 20 µm; (**C** bottom) 10 µm.

### Piezo1 activation inhibits peripheral ruffle formation by inhibition of Rac1 activation

Macropinocytosis is an actin-driven, non-specific endocytosis process accompanied by the following steps: 1) formation of membrane ruffles induced by actin reorganization and 2) subsequent closure of the ruffles to form macropinosomes^36^. We sought to determine which step Yoda1 inhibits. We investigated peripheral ruffle formation by time-lapse live cell imaging and F-actin staining using phalloidin. Peripheral ruffles are actin-rich and sheet-like protrusions of the membrane^37^. In the presence of Yoda1, the extension and folding back of the plasma membrane of A431 cells after EGF addition were clearly observed within 6 min. However, this phenomenon was absent in the presence of Yoda1 (**Figure 2A and Video 1, 2**). In addition, staining actin filaments using phalloidin revealed that Yoda1 inhibited actin polymerization. A431 cells were stimulated with EGF for 5 min in the presence or absence of Yoda1 and then fixed, followed by staining with rhodamine-phalloidin to detect F-actin; then, the cells with the F-actin positive peripheral ruffles were quantified (**Figure 2B**). EGF stimulation resulted in ∼25% of cells with peripheral ruffles, whereas co-treatment with Yoda1 significantly decreased the proportion of cells with peripheral ruffles (∼4%). Moreover, scanning electron microscopy clearly showed that Yoda1 inhibited EGF-induced peripheral ruffle formation (**Figure 2C**).

We then investigated whether Yoda1 inhibited Rac1 activation. In the process of actin rearrangement to form membrane ruffles, EGF-induced actin rearrangement is due to the activation of the small GTPase Rac1^38^. The pulldown experiment of active Rac1 (Rac1-GTP) showed that the amount of Rac1-GTP in the cells treated with EGF increased as previously reported^39^. However, Yoda1 inhibited EGF-induced increase in the amount of Rac1-GTP (**Figure 2D**). These results show that Yoda1 inhibits Rac1 activation.

We examined three possibilities how Yoda1 affected Rac1. First, we investigated whether Yoda1 lowers cytosolic pH. It has been previously shown that macropinocytosis inhibition by amiloride, an inhibitor of Na^+^/H^+^ exchangers (NHE), is due to lower submembranous pH, which prevents Rac1 activation^39^. As previously described, we used the dual-emission ratio (645/585 nm) of seminaphthorhodafluor dye-5 (SNARF-5F) to quantify intracellular pH (**Figure 2— figure supplement 1**). We then compared the cytosolic pH when A431 cells were treated either with dimethyl sulfoxide (DMSO) as vehicle, Yoda1, or ethyl-isopropyl amiloride (EIPA), an amiloride derivative that is widely used as a macropinocytosis inhibitor. EIPA significantly decreased in cytosolic pH, whereas Yoda1 did not lower cytosolic pH (**Figure 2—figure supplement 1B**), suggesting that Rac1 inhibition by Yoda1 is unlikely due to a decrease in intracellular pH. Second, we examined the possibility of inhibition of EGF signal transduction leading to Rac1 activation. Yoda1 did not inhibit EGF-related signaling pathways such as phosphorylation of EGFR and Vav2 (**Figure 2—figure supplement 2**), suggesting that Yoda1 does not affect the acute response of phosphorylation induced by EGF signaling. Finally, we also checked whether Yoda1 affects cholesterol (Chol) distribution in cells. Because membrane ruffling and macropinocytosis in A431 cells require cholesterol to regulate the localization of Rac1^40^, we investigated Chol distribution in the absence or presence of Yoda1 using a genetically encoded biosensor for Chol (mCherry-D4H)^41^. Time-lapse imaging showed that Yoda1 does not affect Chol distribution in the cells, suggesting that inhibition of macropinocytosis by using Yoda1 is not due to change in Chol localization (**Figure 2—figure supplement 3**).

Although the mechanism by which Yoda1 inhibits Rac1 activation remains unclear, these results show that Yoda1 inhibits EGF-induced actin reorganization and peripheral ruffle formation.

### Extracellular Ca^2+^ is required for macropinocytosis inhibition by Piezo1 activation

Since activated Piezo1 is known to be permeable to extracellular Ca^2+^ influx, we next examined whether extracellular Ca^2+^ influx is important for the inhibition of macropinocytosis by Piezo1 activation using Yoda1. To confirm the effects of Yoda1 on Ca^2+^ influx into A431 cells, we conducted time-lapse calcium imaging using A431 cells transiently expressing GCaMP6s. Yoda1 was added 8 min after time-lapse imaging started, resulting in an immediate increase in intracellular Ca^2+^ concentration (**Figure 3A**). Because Piezo1 and other mechanosensitive ion channels could be activated by shear stress such as stimulus by addition of buffer solution itself^27, 42^, Hanks’ balanced salt solution (HBSS) containing DMSO was used as a vehicle control. After adding DMSO solution, the fluorescence of GCaMP6s did not increase, as shown in Fig. 3a, indicating that the intracellular Ca^2+^ concentration did not significantly increase. To investigate whether the intracellular calcium response induced by Yoda1 is due to extracellular Ca^2+^ influx, we used Ca^2+^-free HBSS. Under these conditions, Yoda1 did not increase intracellular Ca^2+^ concentrations (**Figure 3A**). These results indicate that Yoda1 causes extracellular Ca^2+^ influx but does not affect calcium release from intracellular Ca^2+^-storage organelles such as the endoplasmic reticulum.

We then investigated whether Yoda1 inhibits macropinocytosis also in Ca^2+^-free conditions. The dextran uptake assay was conducted using a Ca^2+^-free medium. Yoda1 did not inhibit TMR-dex70 uptake in the absence of extracellular Ca^2+^, indicating that extracellular Ca^2+^ influx through Piezo1 is crucial for the inhibition of macropinocytosis by Yoda1 (**Figure 3B, C**). EGF-induced uptake of TMR-dex70 in Ca^2+^-free medium without Yoda1 was significantly reduced (by ∼40%) compared to that in Ca^2+^-containing medium. EGF-stimulated macropinocytosis in A431 cells has been previously reported to be independent of extracellular Ca^2+^ ion^11^, but the effect of extracellular Ca^2+^ may vary, likely due to differences in experimental conditions and assay systems. However, this result indicates that Yoda1 did not inhibit EGF-stimulated macropinocytosis under extracellular Ca^2+^-free conditions. These data suggest that Piezo1 activation by Yoda1 inhibits macropinocytosis in an extracellular Ca^2+^-dependent manner.

### KCa3.1 activation is necessary for the inhibitory effect of Yoda1 on ruffle formation

We then sought to identify molecule(s) that function downstream of Yoda1-induced Ca^2+^ signaling related to macropinocytosis inhibition. We focused on KCa3.1, a Ca^2+^-activated K^+^ channel that is activated by Ca^2+^ influx through Piezo1 and reduce cell volume in red blood cells^28^. KCa3.1 also plays a key role in EGF-stimulated macropinocytosis^33^. In macropinocytosis, sequential dephosphorylation of phosphoinositides (PI(3,4,5)P_3_ → PI(3,4)P_2_ → PI(3)P → PI) is required^43^. KCa3.1 has been reported to be activated by PI(3)P and is also crucial for macropinocytic cup formation^33, 44^. Therefore, proper temporal activation of KCa3.1 at a later stage of the macropinocytosis process is required for completion of macropincytosis.

We hypothesize that Yoda1-induced Ca^2+^ influx acutely activates KCa3.1, and that the improper activation of KCa3.1 inhibits ruffle formation. Since inhibition of KCa3.1 impairs macropinosome formation but does not affect ruffle formation^33^, the involvement of KCa3.1 in the inhibition of macropinocytosis by Yoda1 was tested by a membrane ruffling assay. A431 cells were pretreated with TRAM-34, a potent and selective KCa3.1 inhibitor, and then treated with EGF and Yoda1 in the presence of TRAM-34. Live cell differential interference contrast (DIC) imaging and phalloidin staining showed that KCa3.1 inhibition by TRAM-34 restored EGF-induced peripheral ruffle formation in the presence of Yoda1 (yellow arrowheads, **Figure 4A, B and Video 3**). We also compared the effects of Yoda1 with ionomycin on the inhibition of macropinocytosis. Ionomycin treatment led to increased intracellular Ca^2+^ concentration (**Figure. 1C**), and it has been reported that ionomycin induces phospholipase C (PLC) activation to hydrolyze PI(4,5)P2 into diacylglycerol (DAG) and inositol-3-phosphate (IP_3_)^45^. PIP_2_ breakdown is thought to lead to inhibition of macropinocytosis, because sequential phosphorylation and dephosphorylation of PIP_2_ is required. Therefore, we checked the amount of PI(4,5)P_2_ in the plasma membrane using a genetically encoded biosensor of PI(4,5)P_2_ (GFP-PLCδ-PH)^46^. Time-lapse imaging showed that ionomycin treatment led to complete redistribution of the PIP_2_ biosensor on the plasma membrane to the cytosol as previously reported^45^, whereas Yoda1 treatment did not induce the redistribution of the biosensor (**Figure 4—figure supplement 1A**). This suggests that Yoda1 treatment, unlike ionomycin, does not hydrolyze PIP_2_ despite increased intracellular Ca^2+^ concentration. In addition, A431 cells pretreated with TRAM-34 did not recover peripheral membrane ruffling in the presence of ionomycin during EGF stimulation (**Figure 4—figure supplement 1B and Video 4**). Altogether, these results suggest that the inhibitory effect of Yoda1 on EGF-stimulated macropinocytosis is related to KCa3.1 activation by an increase in Ca^2+^ concentration through Piezo1 activation; however, the mechanism by which Yoda1 inhibits macropinocytosis is completely different from that of ionomycin.

**Figure 4.**
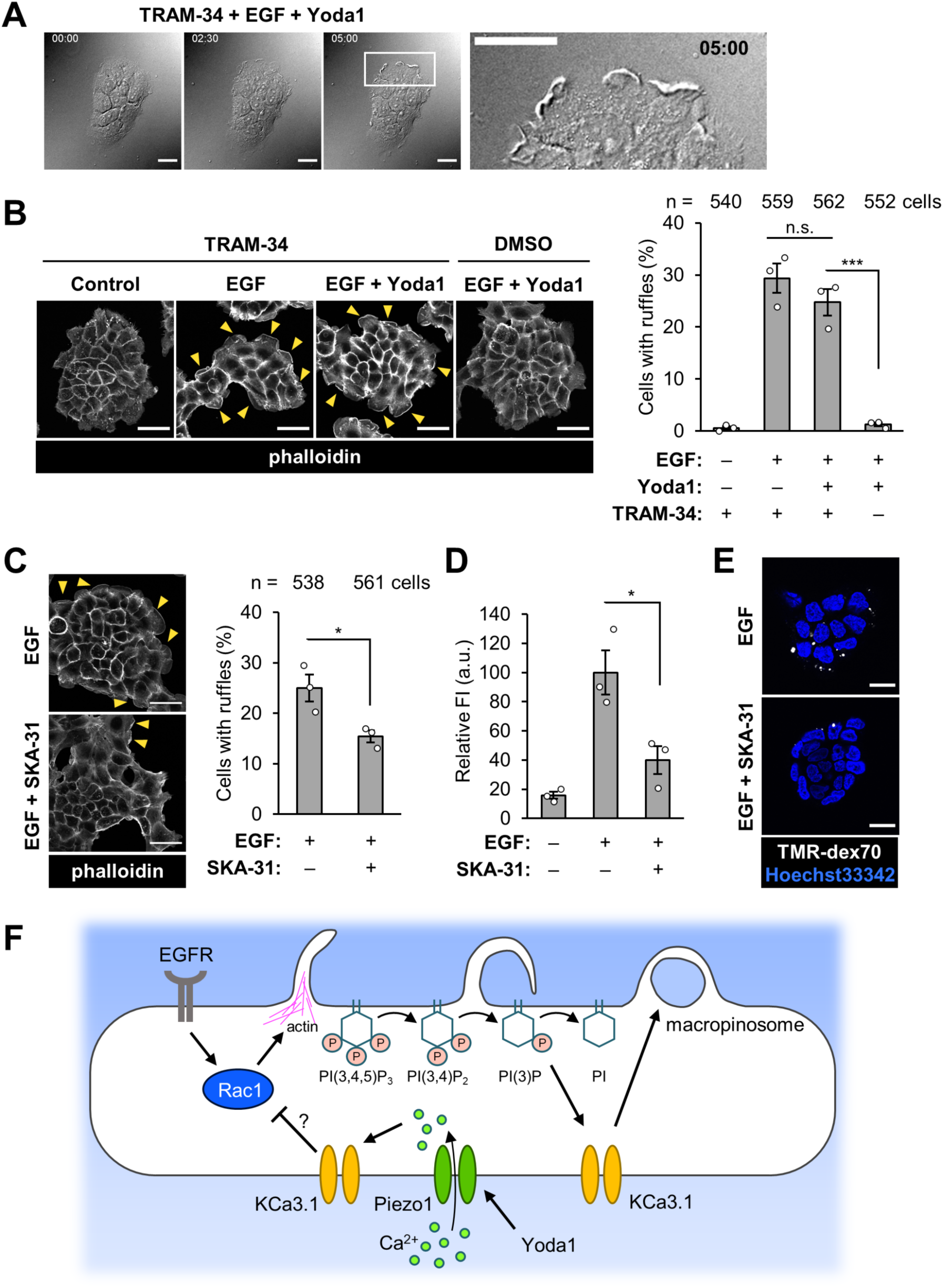
Involvement of KCa3.1 in inhibitory effect of Yoda1 on ruffle formation. (**A**) Live cell imaging of EGF-induced membrane ruffles in A431 cells. The cells were pretreated with a KCa3.1 inhibitor TRAM-34 (10 µM) for 5 min and then stimulated with EGF (20 nM) in the presence of Yoda1 (1.5 µM). Live cell imaging was started immediately after adding EGF and Yoda1. DIC images at the indicated time points (0, 2.5 and 5 min) are shown. The right image shows an enlarged view of the area outlined by the white square in the image at 5 min. (**B**) F-actin staining with phalloidin. Left: Representative images of A431 cells pretreated with TRAM-34 (10 µM) for 5 min and then stimulated with EGF (20 nM) in the absence or presence of Yoda1 (1.5 µM) for 5 min. Right: Quantification of cells with peripheral ruffles. (**C**) F-actin staining with phalloidin. Left: Representative images of A431 cells pretreated with KCa3.1 activator SKA-31 (10 µM) for 5 min and then stimulated with EGF (20 nM) for 5 min. Right: Quantification of cells with peripheral ruffles. (**D**) Flow cytometry analysis of TMR-dex70 uptake. The cells were pretreated with KCa3.1 activator SKA-31 (10 µM) for 5 min and then stimulated with EGF (20 nM) for the uptake of TMR-dex70 for 10 min. (**E**) Confocal microscopy observation of TMR-dex70 uptake. The cells were treated as in (**D**). (**F**) Working hypothesis of macropinocytosis inhibition using Yoda1. Yoda1-induced, Piezo1-dependent extracellular Ca^2+^ influx causes non-proper activation of KCa3.1, which inhibits Rac1 activation followed by ruffle formation. Data in (**B**) and (**C**) represent the mean ± SEM (number of total counted cells pooled from three independent experiments: from left to right, (**B**) 540, 559, 562, and 552; (**C**) 538 and 561). Data in (**D**) represent the mean ± SEM (n = 3 independent biological replicates). *, p < 0.05; ***, p < 0.001; n.s., not significant (one-way ANOVA followed by Tukey-Kramer’s post hoc test (**B**, **D**) or Student’s t-test (**C**)). Yellow arrowheads in (**B**) and (**C**) indicate F-actin positive peripheral membrane ruffling area. Scale bars, (**A**, **E**) 20 µm; (**B**, **C**) 50 µm. **Figure supplement 1.** Yoda1 does not induce PI(4,5)P2 depletion.

### Pharmacological activation of KCa3.1 inhibits EGF-induced macropinocytosis

To further confirm that KCa3.1 activation is involved in the inhibition of macropinocytosis, we next investigated whether the pharmacological activation of KCa3.1 by using SKA-31, a potent potassium channel activator, inhibits EGF-induced membrane ruffle formation and macropinocytic uptake. SKA-31 is known to activate KCa3.1 and KCa2^47^. A431 cells were stimulated with EGF in the absence or presence of SKA-31, and a membrane ruffling assay was performed (**Figure 4C**). Although SKA-31 did not completely inhibit EGF-mediated membrane ruffle induction compared to the effect of Yoda1, there was a significant decrease in the formation of membrane ruffles induced by EGF. Inhibition of macropinocytosis by SKA-31 was also confirmed by flow cytometry analysis and confocal microscopy observation of TMR-dex70 uptake (**Figure 4D, E**). SKA-31 significantly inhibited the uptake of TMR-dex70 by the addition of EGF. Altogether, these results indicate that KCa3.1, if activated in a non-temporal manner, can inhibit EGF-stimulated macropinocytosis.

Taken together with our findings in this study, it is suggested that Yoda1 specifically activates Piezo1, which leads to acute activation of KCa3.1, followed by inhibition of actin rearrangement due to preventing Rac1 activation (**Figure 4F**).

### Piezo1 activation using Yoda1 inhibits cancer cell proliferation

Macropinocytosis functions as a nutrient supply pathway in cancer cells^5, 48^. Therefore, inhibition of macropinocytosis could be a target for a cancer therapy^7, 49^. In this study, we further examined whether Yoda1 treatment for longer incubation (*e.g.*, 16 h) in serum-containing medium inhibits cellular uptake of TMR-dex70 as a marker of macropinocytosis in A431 cells and whether Yoda1 inhibits A431 cell proliferation. A431 cells were incubated with TMR-dex70 and Yoda1 for 16 h, and the cellular uptake of dextran in serum-containing medium was measured by flow cytometry analysis and microscopy observation (**Figure 5A, B**). Even under these experimental conditions, Yoda1 reduced TMR-dex70 uptake in a concentration-dependent manner. TMR-dex70 uptake in 16 h was also inhibited by EIPA, a representative inhibitor of macropinocytosis by inhibiting NHE, suggesting that macropinocytosis may contribute to dextran uptake in A431 cells for 16 h in serum-containing medium (**Figure 5—figure supplement 1**).

**Figure 5.**
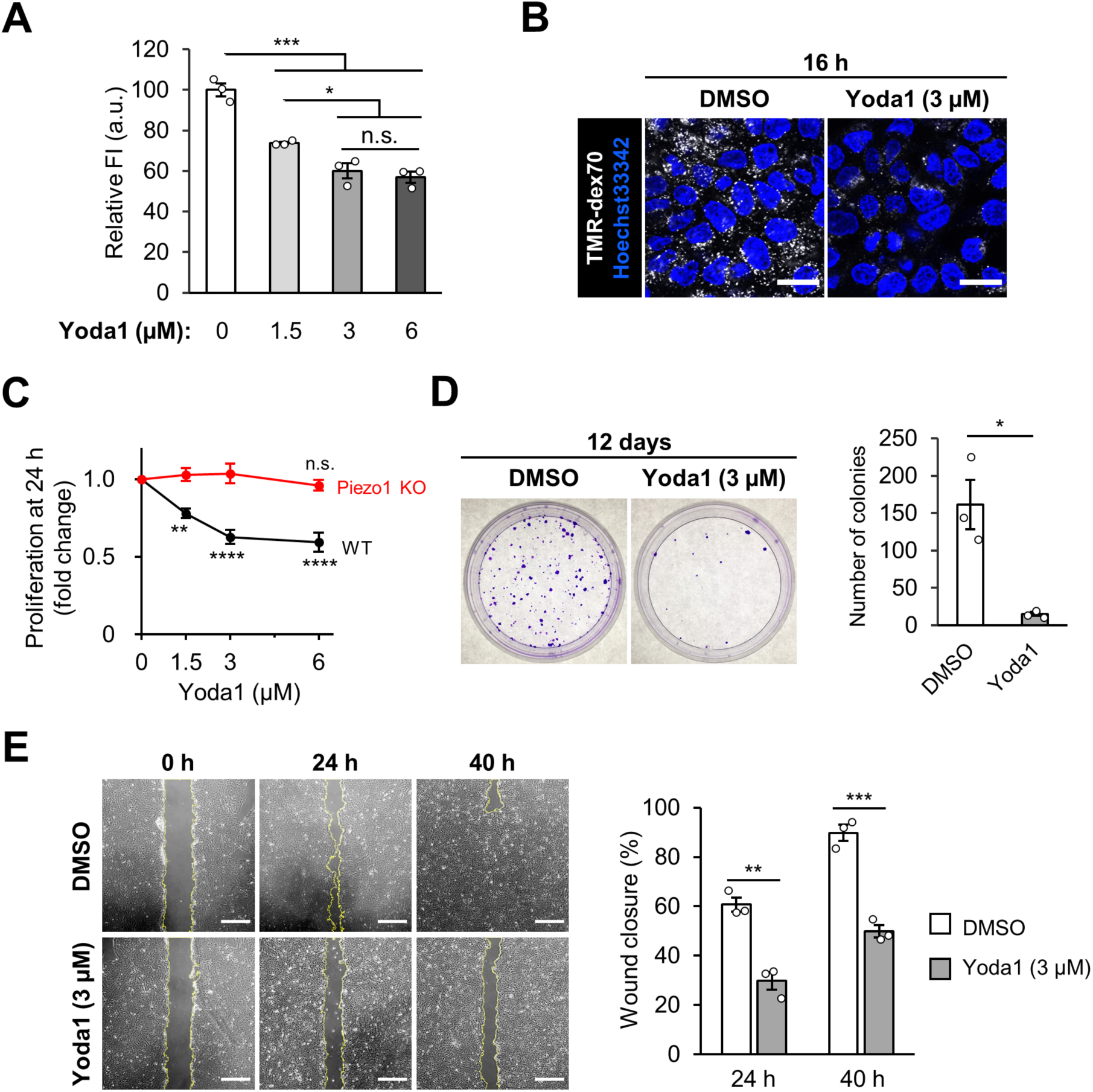
Piezo1 activation inhibits proliferation of A431 cells. (**A**) Flow cytometry analysis of TMR-dex70 uptake into A431 cells treated with Yoda1 at the indicated concentration for 16 h. (**B**) Confocal microscopy observation of TMR-dex70 uptake in 16 h. A431 cells were treated TMR-dex70 in the absence or presence of Yoda1 (3 µM) for 16 h. (**C**) The cell proliferation of A431 cells (WT and Piezo1 KO) in 24 h was investigated by WST-8 assay. The cells were treated with Yoda1 at the indicated concentration for 24 h and then subjected to WST-8 assay. (**D**) The cell survival of A431 cells in the presence of Yoda1 was evaluated by colony formation assay. The cells in 35 mm dishes were treated with DMSO or Yoda1 (3 µM) for 12 days, fixed, stained with 0.2% crystal violet. Left: Representative images. Right: Quantification of the number of visible colonies. Culture medium was changed every 3 days. (**E**) The migration and proliferation of A431 cells in the presence of Yoda1 was measured by wound healing assay. A confluent monolayer of A431 cells was scratched and then treated with DMSO or Yoda1 (3 µM) for 40 h. Left: Representative images. Right: Quantification of the wound closure. Yellow lines indicate the boundary between cells and the scratched area. Data represent the mean ± SEM (n = 3 independent biological replicates) in (**A**), (**C**), (**D**) and (**E**). *, p < 0.05; **, p < 0.01; ***, p < 0.001; ****, p < 0.0001; n.s., not significant (one-way ANOVA followed by Tukey-Kramer’s post hoc test (**A**) or one-way ANOVA followed by Dunnett’s post hoc test (**C**) or Student’s t-test (**D**, **E**)). Scale bars, (**B**) 25 µm; (**E**) 500 µm. **Figure supplement 1.** Inhibition of dextran uptake in 16 h by EIPA and Yoda1. **Figure supplement 2.** Yoda1 inhibits constitutive macropinocytosis and proliferation of HT1080 cells.

A cell proliferation assay using WST-8 formazan, colony formation assay, and wound healing assay were performed to investigate whether Piezo1 activation by Yoda1 inhibited A431 cell proliferation. The WST-8 assay showed that Yoda1 reduced cell proliferation in a concentration-dependent manner (**Figure 5C**). This inhibitory effect was completely abolished in Piezo1 KO A431 cells, demonstrating that decreased proliferation was mediated by Piezo1 activation (**Figure 5C**). Yoda1 treatment at 3 µM showed the maximum effect in these experiments, and as such, we used Yoda1 at 3 µM in the subsequent experiments. The colony formation assay also showed that Yoda1 inhibited long-term cell proliferation (*i.e.*, 12 days) (**Figure 5D**). Furthermore, the wound healing assay indicated that cell proliferation and migration were inhibited by Piezo1 activation using Yoda1 (**Figure 5E**).

We next tested whether Yoda1 has an impact on Ras-transformed cancer cells in which macropinocytosis constantly occurs. Using HT1080 cells, a human fibrosarcoma NRas-mutant cell line, we examined the effect of Yoda1 on dextran uptake and cell proliferation. We confirmed Piezo1 gene expression in HT1080 cells by qPCR (**5—figure supplement 2A**). The dextran uptake assay showed that constitutive macropinocytosis in HT1080 cells was inhibited by Yoda1 in a concentration-dependent manner (**5—figure supplement 2B, C**). In addition, the wound-healing assay showed that proliferation and migration of HT1080 cells were inhibited in the presence of Yoda1, although the effect of inhibition of macropinocytosis and proliferation was not significant compared to that in A431 cells (**5—figure supplement 2D**). However, these data suggest that Yoda1 also has promising activity in the inhibition of constitutive macropinocytosis and cell proliferation in Ras-transformed cells. Altogether, these results suggest that inhibition of macropinocytic cellular uptake through Piezo1 activation leads to the inhibition of cancer cell proliferation.

## Discussion

Our findings demonstrate that Piezo1 activation using Yoda1, a Piezo1 agonist, leads to potent inhibition of EGF-stimulated macropinocytosis in A431 cells. We demonstrated that Yoda1 strongly inhibited the formation of membrane peripheral ruffles induced by EGF. In addition, we also showed that Yoda1 has an inhibitory effect on constitutive macropinocytosis in HT1080 cells, a representative Ras-transformed cancer cell line. In contrast, in Piezo1 KO A431 cells, EGF-stimulated macropinocytosis was observed even in the presence of Yoda1. This result suggests that Piezo1 itself is not crucial for the physiological regulation of macropinocytosis processes that accompany membrane dynamics, such as membrane ruffling. In addition, this study used Yoda1. Although our results cannot strictly conclude that macropinocytosis can be regulated by Piezo1 activity through a physiological increase in membrane tension, we found that Yoda1 inhibits macropinocytosis depending on Piezo1.

We showed that the inhibitory effect of Yoda1 on macropinocytosis was dependent on extracellular Ca^2+^ influx (**Figure 3**). This indicates that Ca^2+^ influx through Piezo1 activation by Yoda1 is essential for the inhibition of macropinocytosis. The inhibition of KCa3.1, which is a calcium-activated potassium channel, recovered the formation of peripheral membrane ruffles in the presence of Yoda1 (**Figure 4A, B**). Therefore, we propose that acute KCa3.1 activation due to Piezo1 activation by Yoda1 likely leads to the inhibition of macropinocytosis. In addition, we showed that a KCa3.1 activator as well as Yoda1 also impaired macropinocytosis (**Figure 4C, D, E**). On the other hand, a previous study reported that KCa3.1 activation is essential to complete macropinosome formation, the later stage of macropinocytosis process^33^. Altogether, our results suggest that appropriate temporal activation of KCa3.1 is important in macropinocytosis.

Furthermore, we demonstrated that Yoda1 treatment inhibited the activation of Rac1, which is essential for membrane ruffling induced by actin rearrangement (**Figure 2D**). A previous study reported that knockdown of Piezo1 in gastric cancer cells led to Rac1 activation^50^. Taken together, our results suggest that Piezo1 activation inhibits Rac1 activation. However, as mentioned above, membrane ruffling was observed even in the presence of Yoda1 in the case of pretreatment with TRAM-34 (**Figure 4A, B**). This result suggests that Rac1 can be activated even in the presence of Yoda1 when KCa3.1 is inhibited. We propose that KCa3.1 activation, following Yoda1-induced Piezo1 activation, could lead to the inhibition of Rac1 activation (**Figure 4F**). Further studies are required to elucidate the relationship between KCa3.1 and Rac1.

Macropinocytosis has recently attracted much more attention, especially from the point of view of cancer metabolism^5, 51, 52^. Preventing macropinocytosis in cancer cells is thought to be an important method for cancer therapy^49^. Therefore, further understanding of the molecular mechanisms and physiological significance of macropinocytosis is required. Unfortunately, there are very few specific and versatile inhibitors for macropinocytosis, because few specific proteins and lipids related to macropinocytosis have been identified^53^. This makes it challenging to develop specific pharmacological tools to inhibit macropinocytosis^54^. Conventional macropinocytosis inhibitors such as cytochalasin D, wortmannin, and EIPA can affect other endocytic pathways or have off-target effects. For instance, cytochalasin D blocks not only macropinocytosis and affects receptor-mediated endocytosis^55^. Wortmannin blocks membrane ruffle closure of macropinocytosis by inhibiting PI3K^56^. Although wortmannin is considered to be a highly selective inhibitor of PI3K, it can also potently inhibit mammalian polo-like kinase 1 (PLK1), which is critical in mitosis^57^. EIPA is one of the most common reagents to inhibit macropinocytosis by blocking NHE, but it also blocks transient receptor potential polycystic 3 (TRPP3), a Ca^2+^-activated channel belonging to the TRP superfamily of cation channels^58^.

This study suggests the possibility of developing a new macropinocytosis inhibitor based on Yoda1. To the best of our knowledge, Yoda1 has so far been considered a highly specific reagent for Piezo1. Although some off-target effects of Yoda1 might exist^59^, our data indicate that Yoda1 inhibits macropinocytosis depending on Piezo1 activation and extracellular Ca^2+^ influx. Interestingly, the inhibitory mechanism of Yoda1 should be clearly different from that of Ca^2+^ elevation using ionomycin, the Ca^2+^ ionophore (**Figure 4A and Figure 4—figure supplement 1A, B**). Thus, the development of novel reagents and methods to increase membrane tension leading to Piezo1 activation might also have the potential to inhibit macropinocytosis.

From the viewpoint of the potential of Yoda1 as a cancer drug, we also demonstrated that Yoda1 has a significant effect on the inhibition of proliferation of A431 cells. This effect is likely dependent on the inhibition of macropinocytosis through Piezo1 activation (**Figure 5B, C, D, E**). These data suggest that Yoda1 and Piezo1 activation may be useful for preventing cancer cell proliferation. In contrast, other studies have suggested that Piezo1 has a key role in promoting cancer cell proliferation and tumor growth^50, 60, 61^. These studies imply that the effect of Piezo1 activation on cell proliferation may vary depending on cell types, and Yoda1 may inhibit the proliferation of specific cancer cells. In addition, methods for targeting cancer cells to activate Yoda1 are required, because Piezo1 has been reported to be involved in various physiological functions^62^, implying that there may be unexpected side effects in normal cells and tissues if Piezo1 is non-selectively activated. Further research is needed to clarify the factors that significantly affect cancer cell proliferation in the presence of Yoda1.

In conclusion, this work is the first to show that the activation of Piezo1 using Yoda1 potently inhibits EGF-stimulated macropinocytosis in A431 cells, which could lead to the inhibition of cell proliferation. Moreover, our results showed that extracellular Ca^2+^ influx through Piezo1 modulates EGF-stimulated macropinocytosis, suggesting the impact of Ca^2+^ on the regulation of macropinocytosis. This study paves the way for the development of novel drugs and methods for studying macropinocytosis study and for cancer therapy by regulating mechanosensitive channel activity.

## Materials and methods

### Reagents

Each reagent was dissolved in the recommended solvent, aliquoted, and stored at −30 °C. Other reagents and culture media were also purchased either from FUJIFILM Wako Pure Chemical Corporation, Sigma-Aldrich, or Thermo Fisher Scientific, unless otherwise specified.

### Cell culture

A431 and HT1080 cells were cultured in Dulbecco’s modified Eagle’s medium (D-MEM, high glucose) (FUJIFILM Wako Pure Chemical Corporation) supplemented with 10% (v/v) heat-inactivated fetal bovine serum (FBS) (Gibco) [D-MEM(+)]. HEK293T cells were cultured in D-MEM with low glucose (FUJIFILM Wako Pure Chemical Corporation) supplemented with 10% (v/v) heat-inactivated FBS. All cells were maintained at 37 °C in a humidified 5% CO_2_ incubator and passaged every 2−4 days. Cells were used for experiments between passage numbers 1 and 15.

### Plasmids construction

pGP-CMV-GCaMP6s-CAAX was a gift from Dr. Tobias Meyer (Addgene, #52228)^63^. pGP-CMV-GCaMP6s was generated by deleting the CAAX sequence from pGP-CMV-GCaMP6s-CAAX (Addgene, #52228). To construct the expression plasmid pIRES2-mCherry, cDNA encoding ZsGreen1 was removed from pIRES2-ZsGreen1 (Takara) by digestion with BstXI/NotI, and then an mCherry cDNA fragment was inserted into the same sites. pPiezo1-IRES2-mCherry was generated by inserting human Piezo1 cDNA (Flexi ORF Clone #FXC01061) into the EcoRI/BamHI sites of pIRES2-mCherry. pPiezo1(Δ946-956)-IRES2-mCherry and pPiezo1(Δ944-946)-IRES2-mCherry were generated by switching from the WT to the deleted sequences between the MluI and SalI sites of pPiezo1-IRES2-mCherry. The deleted sequences were generated as follows: two sequences for each deleted sequence were amplified from pPiezo1-IRES2-mCherry using the following pairs of primers: Piezo1 mutant forward 1 and Piezo1 mutant reverse1, Piezo1 mutant forward2 and Piezo1 mutant reverse2, Piezo1 mutant forward1 and Piezo1 mutant reverse3, and Piezo1 mutant forward3 and Piezo1 mutant reverse2. Then, the two oligos were ligated and digested using the primers forward1 and reverse2 and MluI and SalI, respectively. pSpCas9(BB)-2A-Puro (PX459) V2.0 was a gift from Feng Zhang (Addgene plasmid # 62988; http://n2t.net/addgene:62988; RRID: Addgene_62988)^34^. To construct the CRISPR-Cas9 plasmid for Piezo1-knockout (referred to as PX459-Piezo1), pSpCas9(BB)-2A-Puro (PX459) V2.0 was digested using BbsI-HF (NEB) at 37 °C for 60 min, followed by deactivation at 65 °C for 20 min. After cooling down, QuickCIP (NEB) was added and the mixture was further incubated at 37 °C for 10 min, and then deactivated at 80 °C for 2 min. The digested plasmid was purified using the Wizard SV Gel and PCR Clean-up system (Promega). A guide sequence (TATTCGAGGCCATCGTGTACCGG) to knock out human Piezo1 (accession number: NM_001142864.4) was determined using CRISPRdirect (https://crispr.dbcls.jp). Two oligos, oligo 1 and oligo 2, were phosphorylated using T4 PNK (NEB) and annealed to clone the guide sequence into the sgRNA scaffold of the plasmid. Ligation was then performed by combining the BbsI-digested PX459, the annealed oligo duplex at a 1:3 mol ratio and ligation mix (Takara Bio) at 16 °C for 30 min. The ligation mixture was introduced into *Escherichia coli* DH5α, and the insert sequence was verified by standard sequencing.

### Transfection

Transfection of plasmids was performed using Lipofectamine LTX (Invitrogen) according to the manufacturer’s protocol. Plasmids, Lipofectamine LTX, and PLUS reagent were diluted in Opti-MEM (Invitrogen) and incubated at 25 °C for 5 min for complex formation. The mixture was then added to each dish, resulting in a final plasmid concentration of 2.5 µg/mL. The culture medium was changed 4 h after the transfection. Subsequent experiments were performed 24 h after transfection.

### Establishment of Piezo1-KO A431 cell line

A431 cells (7×10^5^ cells/dish) were seeded onto a 60 mm dish (Iwaki) and incubated overnight. The cells were then transfected with the PX459-Piezo1 plasmid using Lipofectamine LTX (Thermo Fisher Scientific), according to the manufacturer’s instructions, after which they were washed with PBS twice at 6 h after transfection and incubated in D-MEM(+) for 1 day. Afterwards, they were washed with PBS twice and incubated in D-MEM(+) containing puromycin (1 µg/mL) (Sigma) for 3 days. Then the cells were washed twice with PBS and incubated with D-MEM(+) without puromycin for 3 days. Finally, the cells were collected and seeded onto a 96-well plate (Iwaki) by limiting dilution to isolate and culture single-cell clones.

### Sequencing of CRISPR/Cas9 target site of the Piezo1 gene

Genome DNA was extracted from A431 cells (wild type, clones #1 and #2) using GeneArt Genomic Cleavage Detection Kit (Thermo Fisher Scientific), following the manufacturer’s protocol. The sequence around the target of CRISPR/Cas9 was amplified using the following pair of primers: Piezo1 gDNA forward and Piezo1 gDNA reverse. The amplified product was purified using Wizard SV Gel and PCR Clean-up system (Promega) and inserted into T-Vector pMD20 (Takara Bio) using DNA Ligation Kit <Mighty Mix> (Takara Bio) according to the manufacturer’s protocol. The ligation mixture was introduced into *Escherichia coli* DH5α and the insert sequence was verified by standard sequencing.

### RT-PCR and real-time PCR

Total RNA was extracted from A431 cells using NucleoSpin RNA Plus (Takara Bio), following the manufacturer’s protocol. The quantity and quality of RNA was measured by a nanodrop (Thermo Fisher Scientific). RNA concentration was determined by absorbance at 260 nm and RNA quality was confirmed by the 260/280 nm ratio. 2 µg of total RNA was subsequently reverse transcribed to cDNA using PrimeScript RT Master Mix (Takara Bio) according to the manufacturer’s protocol. RT-PCR was done using GoTaq Green Master Mix (Promega). The resulting DNA was fractionated by agarose gel electrophoresis (135 V, 20 min) and viewed with UV transilluminator. Real-time PCR was performed using PowerUp SYBR Green Master Mix (Thermo Fisher Scientific) and 7300 Real-Time PCR System (Applied Biosystems). GAPDH was used as a reference gene.

### Dextran uptake assay

Intracellular uptake of TMR-dex70 was evaluated using confocal microscopy observation and flow cytometry analysis. A431 cells (2×10^5^ cells/dish and 1×10^5^ cells/well) were seeded onto 35 mm glass-bottomed dishes (Iwaki) and a 24-well plate (Iwaki), respectively, and incubated for 1 day. The cells were washed with PBS twice and cultured in D-MEM(−) overnight for serum-starvation. The starved A431 cells were treated with TMR-dex70 (0.5 mg/mL) and reagents as indicated on corresponding figure legends for 10 min at 37 °C. For confocal microscopy observation, the cells were then washed twice with ice-cold PBS and stained nuclei with Hoechst 33342 (5 µg/mL, Invitrogen) for 10 min. The observation was carried out using an FV1000 confocal laser scanning microscope (CLSM) system (Olympus) equipped with a 60× objective lens (UPlanSApo, oil immersion, NA 1.35; Olympus). For flow cytometry analysis, the cells were washed twice with ice-cold PBS, detached from the plate with 0.25% trypsin in PBS for 10 min at 37 °C, added D-MEM(+) to prevent further digestion, and collected into centrifuge tubes. The cells were then centrifuged (800 × *g*, 5 min, 4 °C) and the resulting pellets were washed with ice-cold PBS. The cells were centrifuged again, washed with ice-cold PBS once more and filtrated with a cell strainer. Flow cytometry analysis was performed with 10,000 gated events using an Attune NxT Flow Cytometer (Thermo Fisher Scientific). The results are shown as relative median fluorescence intensity of 10,000 counted events. For HT1080 cells, experiments were conducted without serum starvation. HT1080 cells (2×10^5^ cells/dish and 6×10^4^ cells/well) were seeded onto 35 mm glass-bottomed dishes and a 24-well plate, respectively, and incubated for 1 day. The cells were incubated with TMR-dex70 (0.5 mg/mL) in the absence or presence of Yoda1 in D-MEM(+) for 4 h at 37 °C and then subjected to confocal microscopy observation and flow cytometry analysis as described above.

### Tfn uptake assay

A431 cells (2×10^5^ cells/dish and 1×10^5^ cells/well) were seeded onto 35 mm glass-bottomed dishes and a 24-well plate and incubated for 1 day, and then serum-starved in D-MEM(−) for 1 h prior to experiments. The cells were incubated with AF568-Tfn (20 µg/mL) in the absence or presence of Yoda1 (1.5 µM) in D-MEM(−) for 10 min at 37 °C. The cells were then acid-washed twice with Glycine-HCl buffer (with 150 mM NaCl, pH 3.0) to remove AF568-Tfn on the plasma membrane. Then the cells were fixed with 4% PFA in case of confocal microscopy observation. Microscopy observation and flow cytometry analysis of cellular uptake of Tfn were performed as described above in the method for dextran uptake assay.

### Time-lapse live cell imaging

A431 cells (2×10^5^ cells/dish) were seeded onto 35-mm glass-bottomed dishes (Iwaki) and incubated for 1 day. If necessary, the cells were transfected with the indicated plasmids and serum-starved prior to the experiments. The cells were washed twice with PBS, and the culture medium was replaced with D-MEM(−) (150 µL, on the glass part of the dish). The cells were placed at 37 °C in a microchamber (STXG-IX3WX-SET; Tokai Hit) attached on the FV3000 microscope stage. Reagents in D-MEM(−) (50 µL) were added to the cells to yield the final concentrations indicated in the corresponding figure legends. DIC and fluorescence images were captured every 10 or 20 s using an FV3000 confocal laser scanning microscope (CLSM) system (Olympus) equipped with a 60× objective lens (UPlanSApo, oil immersion, NA 1.35; Olympus).

### Ca^2+^ imaging using GCaMP6s

A431 cells (2×10^5^ cells/dish) seeded onto 35-mm glass-bottomed dishes (Iwaki) were transfected with a plasmid to express GCaMP6s. The cells were washed with Hanks’ balanced salt solution (HBSS; 400 mg/L KCl, 60 mg/L KH_2_PO_4_, 8,000 mg/L NaCl, 350 mg/L NaHCO_3_, 60 mg/L Na_2_HPO_4_ ・ H_2_O, 1,000 mg/L D-Glucose, containing 1 mM Ca^2+^, 1 mM Mg^2+^ and 20 mM HEPES at pH7.4) and the culture medium was replaced with HBSS (150 µL, on the glass part of the dish). GCaMP6s fluorescence images were acquired every 1 min as described above in the method for time-lapse live cell imaging. Fluorescence intensity was measured using ImageJ software (NIH). Yoda1-induced Ca^2+^ influx was quantified as the difference in the GCaMP6s fluorescence intensity between its maximum value (F_max_) and the basal level (F_0_).

### Ca^2+^ imaging using Fluo-8 in HEK293T expressing Piezo1 WT or mutants

HEK293T cells (2×10^5^ cells/dish) were seeded onto 35 mm glass-bottomed dishes and transfected with either Piezo1(full length)-IRES-mCherry, Piezo1(Δ946-956)-IRES-mCherry, Piezo1(Δ944-946)-IRES-mCherry or empty vector (IRES-mCherry). HEK293T cells expressing mCherry were considered to express Piezo1 WT or the mutants. The cells were treated with Fluo-8 AM (5 µM, AAT Bioquest) for 45 min and then washed twice with PBS. The cells were then washed with Hanks’ balanced salt solution (HBSS; 400 mg/L KCl, 60 mg/L KH_2_PO_4_, 8,000 mg/L NaCl, 350 mg/L NaHCO_3_, 60 mg/L Na_2_HPO_4_ ・ H_2_O, 1,000 mg/L D-Glucose, containing 1 mM Ca^2+^, 1 mM Mg^2+^ and 20 mM HEPES at pH7.4) and the culture medium was replaced with HBSS (150 µL, on the glass part of the dish). Fluo-8 fluorescence images were acquired every 1 min as described above in the method for time-lapse live cell imaging. Fluorescence intensity was measured using ImageJ software (NIH). Yoda1-induced Ca^2+^ influx was quantified as the difference in the Fluo-8 fluorescence intensity between its maximum value (F_max_) and the basal level (F_0_).

### Membrane ruffling assay

A431 cells (2.5×10^5^ cells/dish) were seeded onto 35 mm glass-bottomed dishes, cultured for 1 day and then serum-starved overnight prior to experiments. The cells were treated with reagents for 5 min at 37 °C, fixed with 4% paraformaldehyde in PBS for 10 min, and permeabilized with 0.1% Triton X-100 in PBS for 4 min. The cells were then incubated with 1% BSA in PBS for 30 min to block non-specific binding prior to staining F-actin with rhodamine-conjugated phalloidin (Invitrogen) for 30 min. The observations were carried out using the FV1000 CLSM system (Olympus) equipped with a 40× objective lens (UPlanSApo, NA 0.95; Olympus). Among the cells at the margin of the colony, the cells with over 12 µm accumulation of phalloidin were counted as cells with ruffles.

### Scanning electron microscopy

A431 cells were stimulated with EGF (20 nM) in the absence or presence of Yoda1 (1.5 µM) for 5 min and then fixed with 2% glutaraldehyde and 4% paraformaldehyde in 0.1 M cacodylate buffer (pH 7.4) for 2 h, washed with cacodylate buffer, and post-fixed with 1% osmium tetroxide in cacodylate buffer for 2 h. After washing with distilled water, the specimens were subjected to the conductive staining with 1% buffered osmium tetroxide and 1% tannic acid (O-T-O methods). The specimens were then dehydrated in a graded series of ethanol and critical-point drying (Hitachi, Ltd. Critical Point Dryer HCP-1), coated with a thin layer (3 nm) of osmium coater (Vacuum Device Inc.; HPC-30W), and then observed with a field-emission scanning electron microscope (Hitachi S-4800, Tokyo, Japan) at 2 kV acceleration voltage.

### Rac1 activation assay

The Rac1 activation assay was conducted using the Rac1 Pull-Down Activation Assay Biochem Kit (Cytoskeleton) according to the manufacturer’s protocol. A431 cells (9×10^5^ cells/dish) were seeded onto a 100 mm dishes (Iwaki), cultured for 2 days and serum-starved overnight prior to experiments. The cells were treated with reagents for 3 min at 37 °C, lysed with 240 µL of ice-cold lysis buffer [50 mM Tris-HCl (pH 7.5), 10 mM MgCl_2_, 0.5 M NaCl, 2% IGEPAL]. The lysates were cleared by centrifugation at 10,000 × *g* for 1 min at 4 °C. 30 µL of the lysate supernatant was used for protein concentration measurement and preparation of total cell lysate for detecting total Rac1. The remaining supernatant was incubated for 1 h at 4 °C with GST-tagged PAK-PBD beads. After washing the beads, bound proteins were eluted with SDS-loading buffer. The lysates were applied into a polyacrylamide gel (SuperSep Ace, 5-20%, 13well; FUJIFILM Wako Pure Chemical Corporation) and fractionated by SDS-PAGE. The blots were transferred to a polyvinylidene difluoride membrane using Trans-Blot Turbo Transfer System (Bio-Rad), blocked with 5% skim milk in TBS containing 0.05% Tween-20 (TBST) for 1 h at 25 °C, and then incubated overnight at 4 °C with appropriate primary antibodies in 3% skim milk in TBST. After washing the membrane with TBST for 10 min three times, the blots were further incubated with appropriate horseradish peroxidase (HRP)-conjugated secondary antibodies in 3% skim milk in TBST for 1 h at 25 °C. After washing the membrane with TBST for 10 min three times, chemiluminescence was detected using ECL prime (GE Healthcare) and LAS3000 mini (FUJIFILM). The images were analyzed using ImageJ software (NIH).

### Western blot

A431 cells (3×10^5^ cells/well) were seeded on a 6-well plate (Iwaki), cultured for 1 day and serum-starved overnight prior to experiments. The cells were treated with EGF in the presence or absence of Yoda1 for 5 min at 37 °C, lysed with ice-cold RIPA buffer [50 mM Tris-HCl (pH 7.6), 150 mM NaCl, 1 mM EDTA, 1% Triton X-100, 0.1% SDS, 0.1% sodium deoxycholate] containing protease inhibitor cocktails (Roche) and phosphatase inhibitor cocktails (Roche), and lysates were cleared by centrifugation at 16000 × *g* for 20 min at 4 °C. The protein concentrations of the lysates were measured by BCA protein assay using Pierce BCA Protein Assay Kit (Thermo Fisher Scientific) and then unified to 0.5 µg/µL. SDS-PAGE, antibody treatment and detection were performed described as above in the method for Rac1 activation assay using 5% BSA in TBST as blocking solution. When detecting EGFR and Vav2, the membrane was subjected for the first immunoblots (pEGFR or pVav2), stripped by immerging the membrane in Restore PLUS Western Blot Stripping Buffer (Thermo Fischer Scientific) for 5 min and washed twice with TBST for 10 min and then subjected to the second immunoblots (EGFR or Vav2).

### Intracellular pH measurement

A431 (2.5×10^5^ cells/dish) cells were seeded on 35 mm glass-bottomed dishes (Iwaki) and incubated for 1 day. The cells were incubated with SNARF-5F AM (20 µM, Invitrogen) in D-MEM(−) for 30 min at 37 °C, washed twice with PBS, and then incubated with Yoda1 (1.5 µM) in D-MEM(−) for 10 min at 37 °C. EIPA (25 µM, 30 min, 37 °C) was used as a positive control which decreases intracellular pH. The observation was done using the FV1000 CLSM system (Olympus) equipped with a 40× objective lens (UPlanSApo, NA 0.95; Olympus). SNARF-5F AM was excited at 559 nm and images were acquired in the range of 570-600 nm and 630-660 nm to evaluate emission signal at 585 nm and 645 nm, respectively. Fluorescence intensity was measured using ImageJ software (NIH) and ratio of emission signal of 585/645 nm was calculated. For establishing calibration curve, cells were incubated with SNARF-5F AM as above, washed twice with calibration buffer (130 mM KCl, 10 mM NaCl, 1 mM MgSO_4_, 10 mM MOPS) at pH 6.2, 6.6, 7.0, 7.4, 7.8, or 8.2, and then incubated with calibration buffer containing 10 µg/mL nigericin for 15 min at 25 °C to equilibrate the intracellular pH with extracellular pH.

### WST-8 assay

The WST-8 assay was performed using the Cell Counting Kit-8 (Dojindo), according to the manufacturer’s protocol. A431 cells (1×10^4^ cells/well) were seeded onto a 96-well plate (Iwaki) and incubated for 1 day. The cells were treated with DMSO or Yoda1 (1.5, 3, or 6 µM) in D-MEM(+) for 24 h at 37 °C. The cells were then washed twice with PBS before adding 100 µL D-MEM(−) and 10 µL of WST-8 reagent to each well. The cells were further incubated for 2 h at 37 °C followed by measuring the absorbance at 450 nm.

### Colony formation assay

A431 cells (4×10^2^ cells/dish) were seeded onto 35 mm dishes and incubated for 1 day. The cells were treated with DMSO or Yoda1 (3 µM) in D-MEM(+) for 12 days at 37 °C, until the DMSO-treated cells formed visible colonies. The culture medium was changed every 3 days. The cells were then fixed with 4% paraformaldehyde for 20 min and stained with 0.2% crystal violet for 20 min. The dishes were photographed, and visible colonies were counted.

### Wound healing assay

A431 cells (4×10^5^ cells/dish) were seeded onto 35 mm dishes and incubated for 2 days. A confluent monolayer of A431 cells was scratched with a 10 μL pipette tip to create a scratch wound. The cells were then treated with DMSO or Yoda1 (3 µM) in D-MEM(+) for 40 h at 37 °C. Phase contrast images of the wounded areas at 0, 24 and 40 h after scratching were acquired using a CK40 inverted microscope (Olympus) equipped with a 4× objective lens (SPlan 4PL, NA 0.13; Olympus) and a 10× eyepiece lens (NCWHK; Olympus). The wounded areas were measured using ImageJ software (NIH).

### Statistical analysis

No statistical method was used to determine the sample size prior to the study. All data are presented as the mean ± standard error of the mean (SEM) of three independent biological replicates (n = 3) unless otherwise specified. All statistical analyses were performed using JMP Pro 14. For comparison of two groups, an unpaired Student’s t-test was used. For multiple comparison analyses, one-way analysis of variance (ANOVA) followed by Tukey-Kramer’s post hoc test or Dunnett’s post hoc test was used. The calculated P-values were considered significant at P < 0.05.

## Supporting information

Key Resource Table

## Acknowledgements

pGFP-C1-PLCδ-PH and pmCherry-C1-D4H were kind gifts from Dr. Gregory Fairn (St. Michael’s Hospital, Toronto, Canada)^41^. This work was supported by JST CREST (Grant Number JPMJCR18H5).

## Author contributions

H.H. conceived the project, designed the experiments, established the Piezo1 knockout A431 cells with the help of Y.H., and analyzed the data. M.K. designed and performed most experiments and analyzed the data. T.M., J.V.V.A., and M.I. contributed to analysis for flow cytometry, Rac1 pulldown assay, and plasmid construction for Piezo1 mutants, respectively. M.S. and M.M. performed the scanning electron microscopy observations. Y.H. contributed to construction of Piezo1-related plasmids and establishment of the Piezo1 knockout cell line. M.K. and H.H. wrote the original manuscript. S.F. supervised the project and wrote the manuscript with H.H. and M.K.. All authors discussed and commented on the manuscript.

## Supporting Information

**Figure 1—figure supplement 1.**
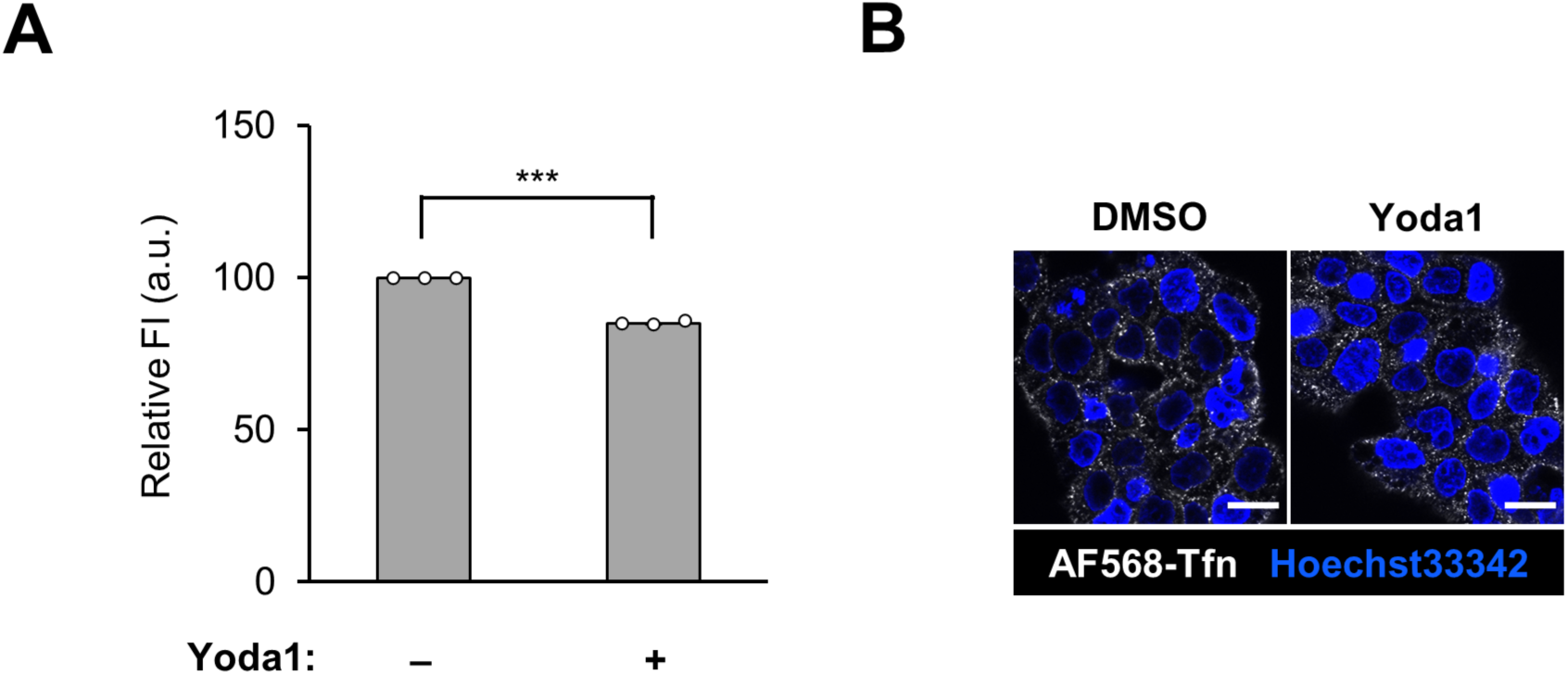
Tfn uptake in the absence or presence of Yoda1. (**A**) Flow cytometry analysis of AF568-Tfn uptake into A431 cells. The cells were incubated with AF568-Tfn (20 µg/mL) in the absence or presence of Yoda1 (1.5 µM) for 10 min. Data represent the mean ± SEM (n = 3 independent biological replicates). (**B**) Confocal microscopy observation of AF568-Tfn uptake. A431 cells were treated as (**A**). Scale bars, 20 µm.

**Figure 1—figure supplement 2.**
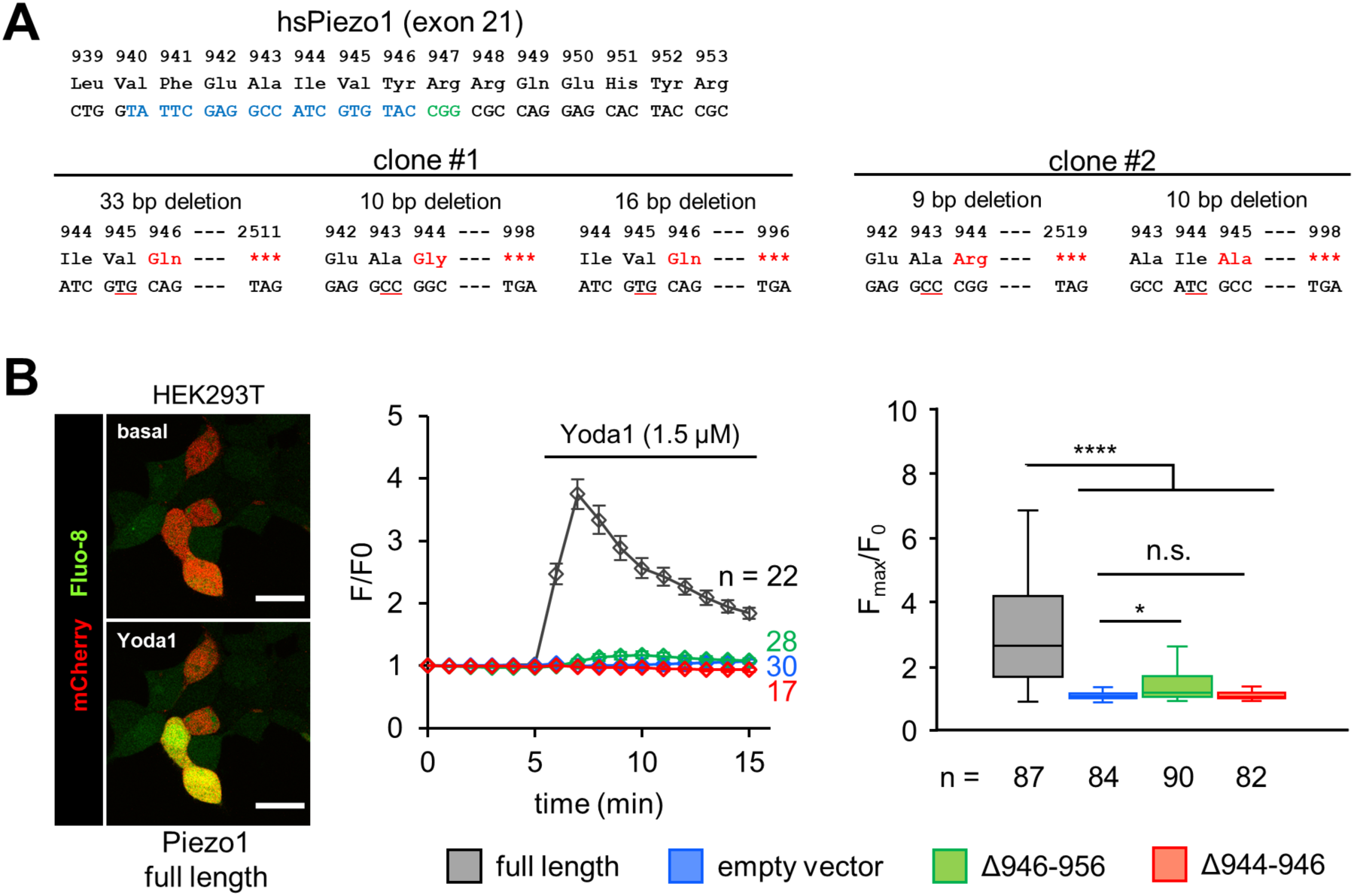
Supporting data for characterization of Piezo1 knockout A431 cells. (**A**) CRISPR/Cas9 target site of the Piezo1 gene. The guide sequence and the protospacer adjacent motif are showed in blue and green characters, respectively. CRISPR/Cas9-mediated deletions are showed in red characters. (**B**) HEK293T cells were transfected with indicated vector, loaded with Fluo-8 AM (5 µM) and then Fluo-8 fluorescence intensity was recorded every 1 min. Transient gene expression was confirmed by IRES2 dependent co-expression of mCherry. Yoda1 (1.5 µM) was added 5 min after start of time-lapse imaging. Left: Representative images before and after Yoda1 addition. Middle: Representative time-course of relative fluorescence intensity of Fluo-8 AM. Data represent the mean ± SEM. Right: Quantification of maximum Yoda1-induced GCaMP6s intensity increase. (n ≥ 82 cells for each condition pooled from three independent experiments.) Box and whiskers graph: line, median; box, upper and lower quartiles; whiskers, maxima and minima; circle plots, outliers. Scale bars, 25 µm. ****, P < 0.0001; *, P < 0.05; n.s., not signficant (one-way ANOVA followed by Tukey– Kramer’s post hoc test).

**Figure 1—figure supplement 3.**
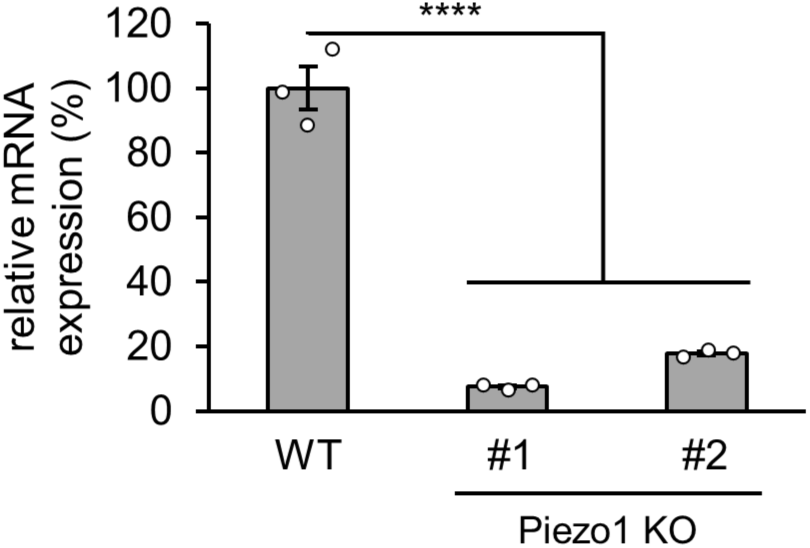
Piezo1 gene expression examined by qPCR. Expression of Piezo1 in A431 cells (WT, KO clone #1, 2). Piezo1 mRNA expression level was determined by qPCR. The GAPDH shows relative mRNA expression level of Piezo1 in each condition. Data represent mean ± SD of triplicate samples from one of two independent experiments. ****, P < 0.0001 (one-way ANOVA followed by Dunnett’s post hoc test).

**Figure 2—figure supplement 1.**
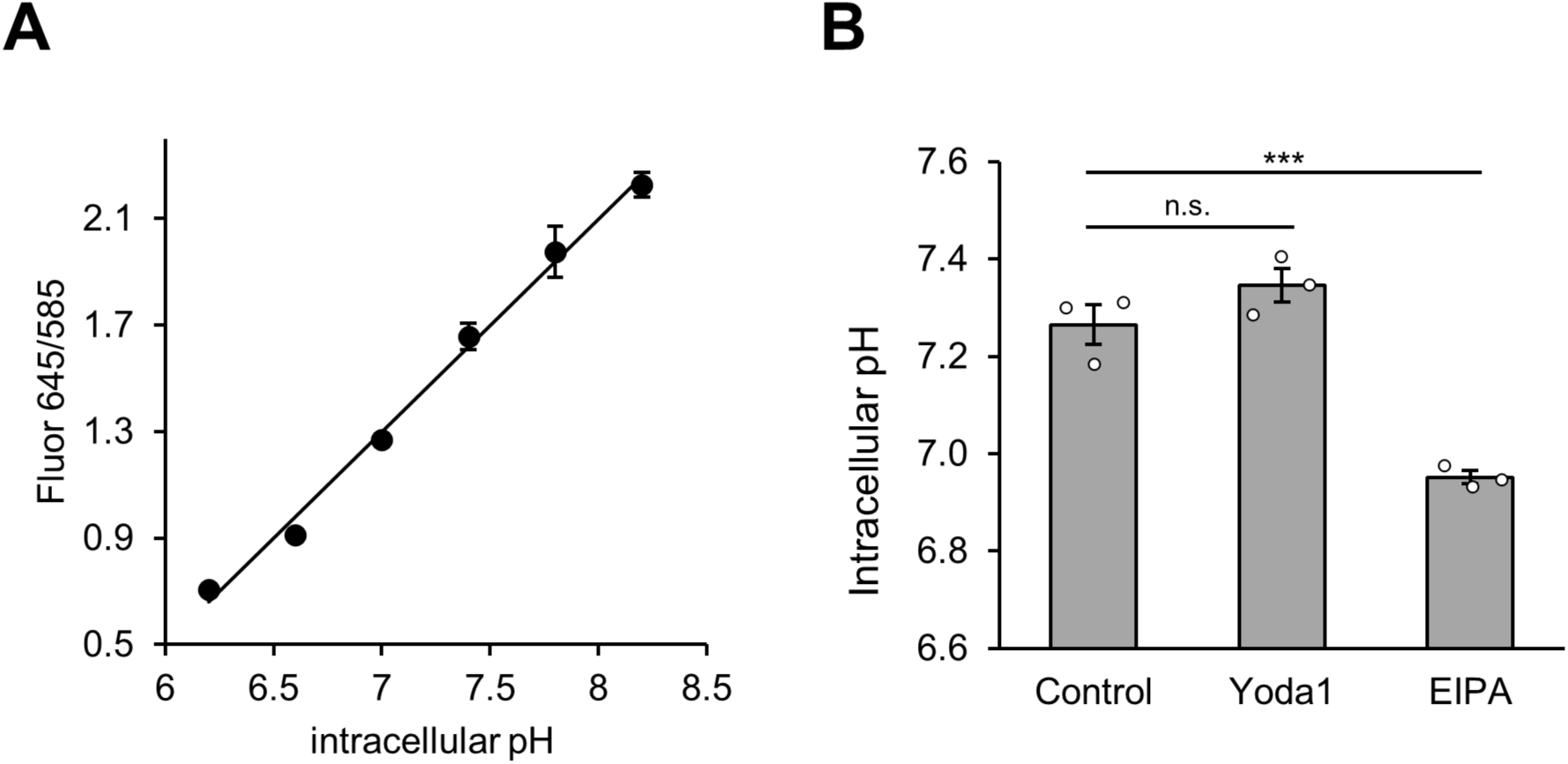
Yoda1 does not change intracellular pH. (**A**) Relationship between fluorescence intensity ratio (645/585 nm) of SNARF-5F AM and intracellular pH. Fluorescence intensity ratio (645/585 nm) of SNARF-5F AM was plotted against respective pH and linear regression analysis was performed to obtain the calibration curve. Data represent mean ± SEM (n = 150 cells for each condition from three independent experiments). (**B**) Intracellular pH measurement using SNARF-5F AM in starved A431 cells treated either with Yoda1 (1.5 µM) for 10 min or EIPA (25 µM) for 30 min in D-MEM(–). EIPA was used as a positive control to reduce intracellular pH. The intracellular pH value was obtained from the calibration curve in (**A**). Data represent mean ± SEM (n = 150 cells for each condition from three independent experiments). ***, P < 0.001; n.s., not significant (one-way ANOVA followed by Dunnett’s post hoc test).

**Figure 2—figure supplement 2.**
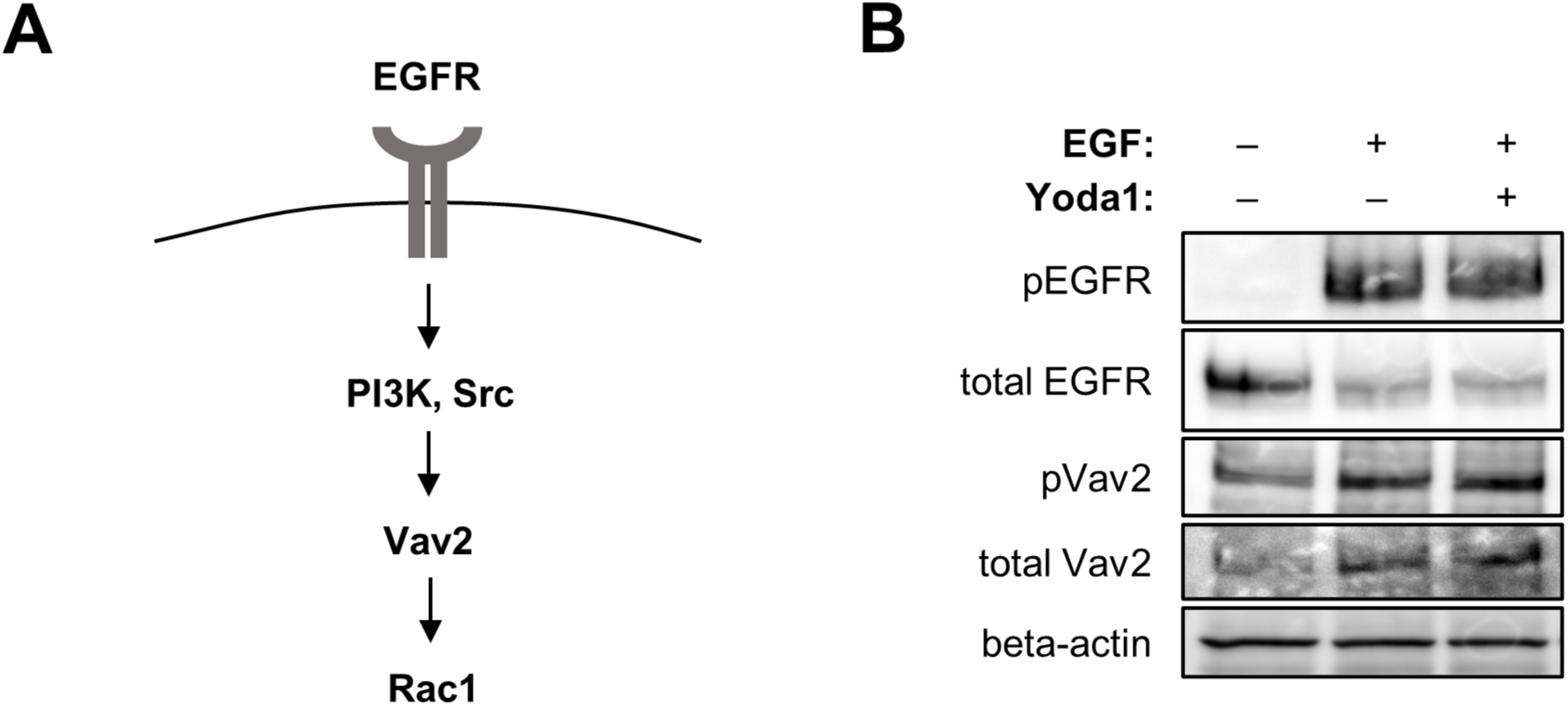
Yoda1 does not inhibit phosphorylation of EGFR and Vav2. (**A**) Schematic diagram showing the pathway of Rac1 activation by EGF. (**B**) Western blot analysis. A431 cells were treated with EGF (20 nM) and Yoda1 (1.5 μM) for 5 min, and total cell lysates were analyzed by performing SDS-PAGE followed by western blot. The experiments were performed at least twice with similar results.

**Figure 2—figure supplement 3.**
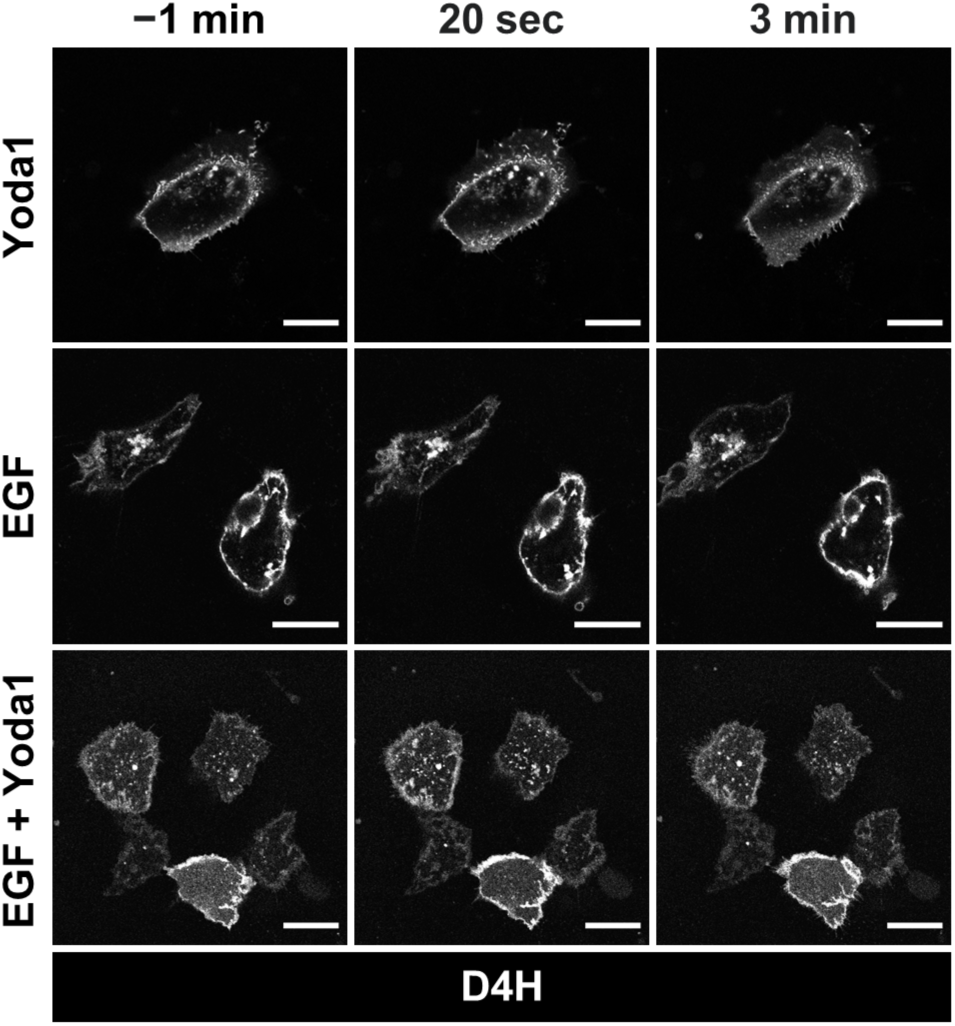
Yoda1 does not affect cholesterol localization. A431 cells were transfected with mCherry-tagged D4H, a specific probe for cholesterol, serum-starved for 4 h, and then mCherry-D4H fluorescence images were acquired every 20 sec. The time of EGF (20 nM) and Yoda1 (1.5 µM) addition was defined as time = 0. Images at indicated time points (1 min before adding reagents and 20 sec, 3 min after addition) are shown. Scale bars, 20 µm.

**Figure 4—figure supplement 1.**
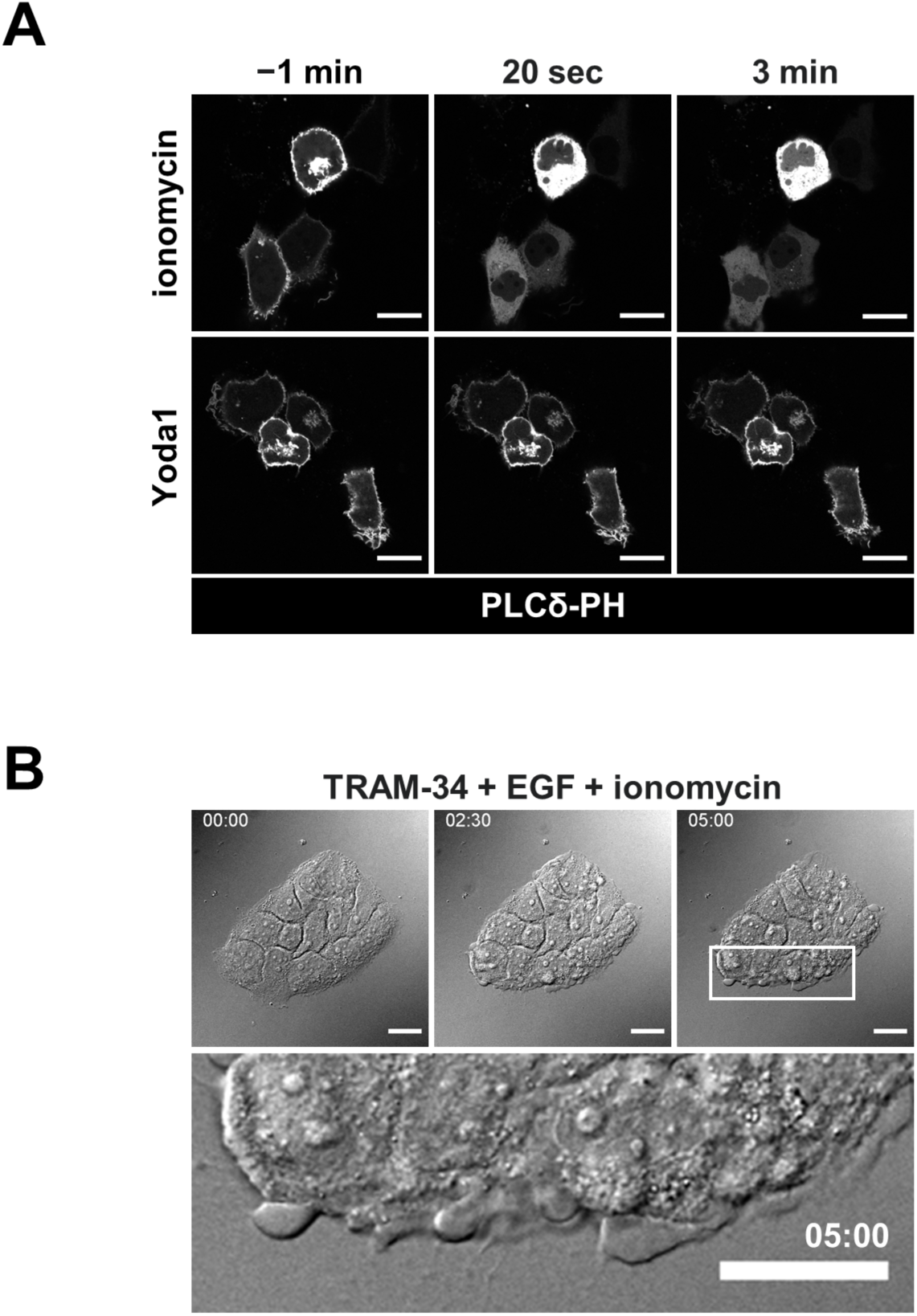
Yoda1 does not induce PI(4,5)P2 depletion. (**A**) A431 cells were transfected with GFP-tagged PLCδ-PH, a specific probe for PI(4,5)P2, and then GFP-PLC-PH fluorescence images were acquired every 20 s. The time of ionomycin (5 µM) or Yoda1 (1.5 µM) addition was defined as time = 0. Images at indicated time points (1 min before adding reagents, 20 s and 3 min after addition) are shown. (**B**) Live imaging shows that ionomycin inhibits EGF-induced ruffle also in the presence of TRAM-34. A431 cells were pretreated with a KCa3.1 inhibitor TRAM-34 (10 µM) for 5 min and then stimulated with EGF (20 nM) in the presence of ionomycin (5 µM). Time-lapse imaging was started immediately after adding EGF and ionomycin. DIC images at the indicated time points (0 min, 2.5 min and 5 min) are shown. The bottom image shows an enlarged view of the area outlined by the white square in the image at 5 min. Scale bars, (**A**, **B**) 20 µm.

**Figure 5—figure supplement 1.**
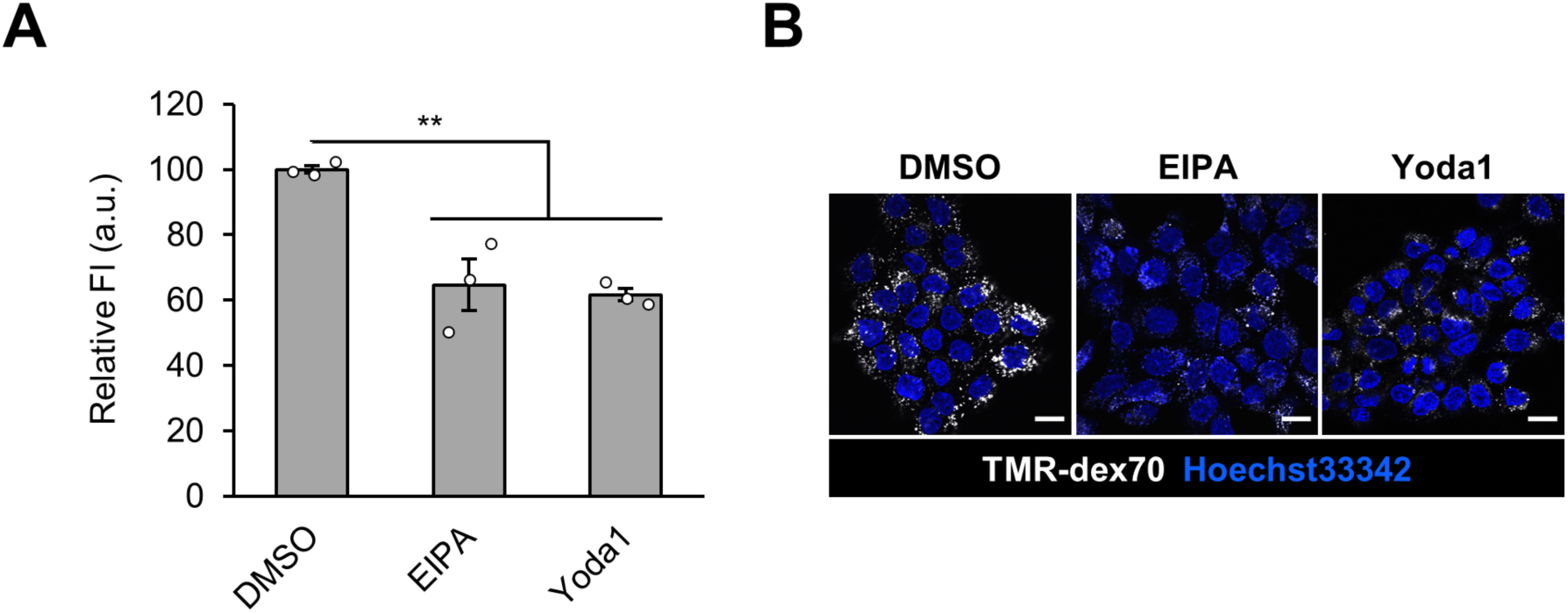
Inhibition of dextran uptake in 16 h by EIPA and Yoda1. (**A**) Flow cytometry analysis of TMR-dex70 uptake into A431 cells. The cells were treated with TMR-dex70 (0.5 mg/mL) in the presence of DMSO as vehicle control, EIPA (20 µM) or Yoda1 (1.5 µM) in serum-containing medium for 16 h. (**B**) Confocal microscopy observation of TMR-dex70 uptake in 16 h. A431 cells were treated as (**A**). Data represent the mean ± SEM (n = 3 independent biological replicates) in (**A**). **, p < 0.01 (one-way ANOVA followed by Dunnett’s post hoc test). Scale bars, 20 µm.

**Figure 5—figure supplement 2.**
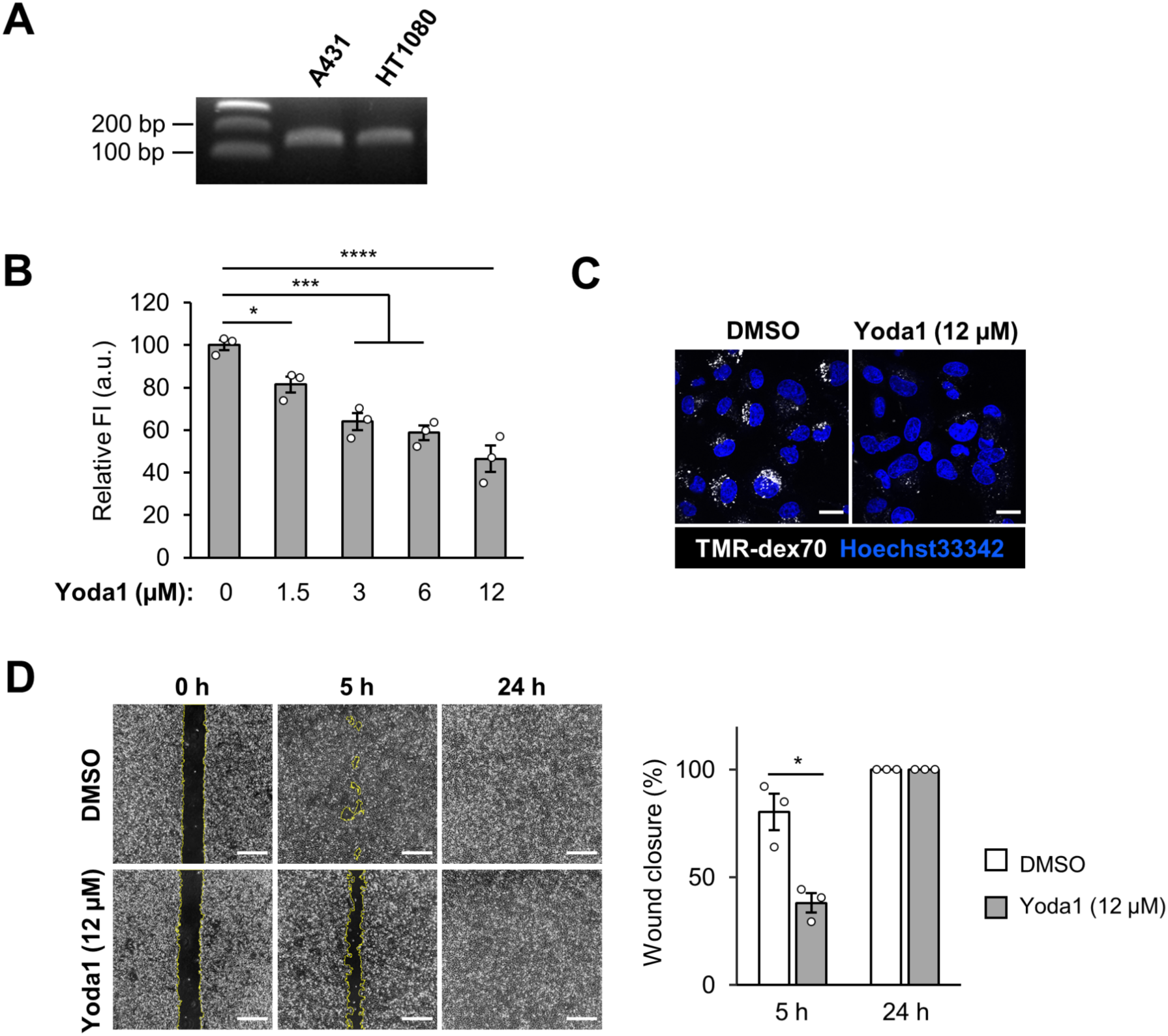
Yoda1 inhibits constitutive macropinocytosis and proliferation of HT1080 cells. (**A**) Expression of Piezo1 in HT1080 cells. mRNA expression of Piezo1 was confirmed by RT-PCR. (**B**) Flow cytometry analysis of TMR-dex70 uptake into A431 cells. The cells were incubated with TMR-dex70 (0.5 mg/mL) in the absence or presence of Yoda1 at indicated concentration for 4 h. (**C**) Microscopy observation of TMR-dex70 uptake. The cells were incubated with TMR-dex70 (0.5 mg/mL) in the absence or presence of Yoda1 (12 µM) for 4 h. (**D**) The cell migration and proliferation of HT1080 cells in the presence of Yoda1 were measured by wound healing assay. A confluent monolayer of HT1080 cells was scratched, treated with DMSO or Yoda1 (12 µM) for 24 h. Left: Representative images. Right: Quantification of the wound closure. *, p < 0.05; ***, p < 0.001; ****, p < 0.0001 (one-way ANOVA followed by Dunnett’s post hoc test (**C**)). Scale bars, (**C**) 20 µm; (**D** right) 500 µm.

## Videos

**Video 1.** EGF induces peripheral ruffle formation. Differential interference contrast (DIC) movie of A431 cells treated with EGF (20 nM). Time-lapse imaging was started immediately after adding EGF, and frames were acquired every 10 s. Video was sped up 100× (ten frames per second) over real time. Scale bar, 20 μm.

**Video 2.** Yoda1 inhibits ruffle peripheral formation. Differential interference contrast (DIC) movie of A431 cells treated with EGF (20 nM) and Yoda1 (1.5 μM). Time-lapse imaging was started immediately after adding EGF and Yoda1, and frames were acquired every 10 s. Video was sped up 100× (ten frames per second) over real time. Scale bar, 20 μm.

**Video 3.** TRAM-34 recovers peripheral formation in the presence of Yoda1. Differential interference contrast (DIC) movie of A431 cells pretreated with TRAM-34 (10 μM) for 5 min and then treated with EGF (20 nM) and Yoda1 (1.5 μM). Time-lapse imaging was started immediately after adding EGF and Yoda1, and frames were acquired every 10 s. Video was sped up 100× (ten frames per second) over real time. Scale bar, 20 μm.

**Video 4.** TRAM-34 does not recover peripheral formation in the presence of ionomycin. Differential interference contrast (DIC) movie of A431 cells pretreated with TRAM-34 (10 μM) for 5 min and then treated with EGF (20 nM) and ionomycin (5 μM). Time-lapse imaging was started immediately after adding EGF and ionomycin, and frames were acquired every 10 s. Video was sped up 100× (ten frames per second) over real time. Scale bar, 20 μm.

